# Divergent acute *versus* prolonged pharmacological GLP-1R responses in adult beta cell-selective β-arrestin 2 knockout mice

**DOI:** 10.1101/2022.04.21.489075

**Authors:** Stavroula Bitsi, Yusman Manchanda, Liliane ElEid, Nimco Mohamed, Ben Hansen, Kinga Suba, Guy A. Rutter, Victoria Salem, Ben Jones, Alejandra Tomas

**Affiliations:** Section of Cell Biology and Functional Genomics, Division of Diabetes, Endocrinology and Metabolism, Department of Metabolism, Digestion and Reproduction, Imperial College London, London, UK; Department of Bioengineering, Imperial College London, London, UK; CHUM Research Centre, University of Montreal, Quebec, H2X 0A9, Canada; Lee Kong Chian School of Medicine, Nanyang Technological University, 637553, Singapore; Section of Endocrinology and Investigative Medicine, Division of Diabetes, Endocrinology and Metabolism, Department of Metabolism, Digestion and Reproduction, Imperial College London, London, UK

## Abstract

The glucagon-like peptide-1 receptor (GLP-1R) is a major therapeutic target in type 2 diabetes (T2D) and obesity. Following activation, GLP-1Rs are rapidly desensitised by β-arrestins, scaffolding proteins that terminate G protein interactions but also act as independent signalling mediators. While GLP-1R interacts with β-arrestins 1 and 2, expression of the latter is greatly enhanced in beta cells, making this the most relevant isoform. Here, we have assessed *in vivo* glycaemic responses to the pharmacological GLP-1R agonist exendin-4 in adult beta cell-selective β-arrestin 2 knockout (KO) mice. Lean female and high-fat, high-sucrose-fed KO mice of both sexes displayed worse acute responses *versus* control littermates, an effect that was inverted 6 hours post-agonist injection, resulting in prolonged *in vivo* cell-cell connectivity in KO islets implanted in mouse eyes. Similar effects were observed for the clinically relevant semaglutide and tirzepatide but not with exendin-phe1, an agonist biased away from β-arrestin recruitment. *Ex vivo* acute cAMP was impaired, but overnight desensitisation was reduced in KO islets. The acute signalling defect was attributed to enhanced β-arrestin 1 and phosphodiesterase (PDE) 4 activity in the absence of β-arrestin 2, while the reduced desensitisation correlated with altered GLP-1R trafficking, involving impaired recycling and lysosomal targeting and increased trans-Golgi network (TGN) localisation and signalling, as well as reduced GLP-1R ubiquitination by the E3 ubiquitin ligase NEDD4. This study has unveiled fundamental aspects of the role of β-arrestin 2 in regulating pharmacological GLP-1R responses with direct application to the rational design of improved GLP-1R-targeting therapeutics.

## Introduction

The glucagon-like peptide-1 receptor (GLP-1R), a class B G protein-coupled receptor (GPCR), is a prominent target for type 2 diabetes (T2D) and obesity treatment. Binding of endogenous GLP-1 to beta cell GLP-1Rs potentiates postprandial insulin secretion, with pharmacological agonists successfully leveraging this effect to control blood glucose levels in people with T2D (1). GLP-1R agonists are nonetheless associated with dose-related side effects such as nausea and diarrhoea, negatively impacting tolerability and reducing the range of acceptable dosages (2). Exploiting pathway selectivity downstream of GLP-1R activation to potentiate beneficial over detrimental responses has been proposed as a strategy to increase effectiveness, reduce side effects, and improve adherence (3–5).

The arrestins, a family of cytosolic adaptor proteins, were first identified as key contributors to GPCR homologous desensitisation, thereby ‘arresting’ GPCR signalling (6, 7). Later research assigned them an additional role as *bona fide* signalling mediators (8, 9), contributing to the concept of ‘biased agonism’, whereby either G protein or arrestin-mediated pathways are preferentially activated. In contrast to arrestins 1 and 4, whose expression is confined to the visual system, β-arrestins 1 and 2 (also known as arrestins 2 and 3, respectively) are ubiquitously expressed (10). β-arrestins are classically associated with receptor internalisation via clathrin-coated pits (11, 12); however, we and others have shown that they are dispensable for GLP-1R endocytosis (13–15). Conversely, there is building evidence that the interaction of GLP-1R and β-arrestins results in autonomous signalling events (14, 16). Interestingly, GLP-1R pharmacological agonists biased away from β-arrestin recruitment exhibit improvements in signalling duration and capacity for sustained insulin secretion (5), while agonist-induced cyclic AMP (cAMP) / protein kinase A (PKA) signalling is prolonged in β-arrestin 1/2 double-knockout (KO) HEK293 cells (17), suggesting that β-arrestin deficiency favours sustained GLP-1R action.

The two β-arrestin isoforms result from alternative mRNA splicing (18) and, despite sharing structural and functional similarities, often exhibit differential and at times contrasting actions (19–21). Importantly, β-arrestin 2 is much more abundant than β-arrestin 1 in pancreatic islets, with ∼50-fold increase in mRNA levels in both human and mouse beta cells (22, 23). Whole-body constitutive (24) and beta cell-selective tamoxifen-inducible β-arrestin 2 KO (25) murine models display compromised glucose-stimulated insulin secretion and glucose tolerance on high-fat but not regular chow diet. The latter study attributed these impairments to calcium/calmodulin-dependent protein kinase II (CaMKII)-dependent mechanisms. However, the effect of β-arrestin 2 deletion in the control of pharmacological GLP-1R responses was not investigated *in vivo* in these mouse models, and only one *ex vivo* experiment in islets extracted from lean beta cell-specific β-arrestin 2 KO male mice was reported, suggesting no significant alterations in acute calcium signalling or insulin secretion under these conditions (25).

Thus, although knockout studies suggest that loss of β-arrestin 2 results in deleterious effects on whole-body glucose metabolism (particularly under metabolic stress), pharmacological manipulation of GLP-1R agonists away from recruitment of both β-arrestins seems to improve their glucose-lowering potency, particularly over prolonged periods. Additional discrepancies persist, such as the conflicting reports on the effect of β-arrestin on GLP-1R internalisation. Overall, the role of β-arrestin 2 in modulating pharmacological GLP-1R responses in primary beta cells, and its associated mechanisms, remain poorly characterised, especially *in vivo*.

Given the potential significance of β-arrestin 2 in modulating pancreatic GLP-1R function, we have determined here both acute and sustained pharmacological GLP-1R responses by using a range of *in vivo*, *ex vivo* and *in vitro* approaches and mouse models, unveiling a previously unknown dual role of β-arrestin 2 in the control of GLP-1R signalling, with an initial acute potentiating effect that then progresses to a dampening effect over prolonged agonist stimulation periods. We have additionally identified a compensatory activity of β-arrestin 1 and the cAMP phosphodiesterase PDE4 as being responsible for the acute GLP-1R signalling defect detected in β-arrestin 2-deleted beta cells. Finally, we have unveiled changes in GLP-1R post-endocytic trafficking and ubiquitination signatures, which correlate with altered GLP-1R association with the ubiquitin ligase NEDD4 and GLP-1R signal prolongation in the absence of β-arrestin 2.

## Results

### Deletion of beta cell β-arrestin 2 exerts sex- and dose-dependent effects on GLP-1R agonism in vivo

To explore the importance of β-arrestin 2 in the pharmacological GLP-1R responses from adult animals, we generated a beta cell-selective, tamoxifen-inducible β-arrestin 2 KO mouse model (Figure 1A, Supplemental Figure 1A; Pdx1-Cre-ERT/Barr2 fl/fl and control Barr2 fl/fl mice). β-arrestin 2 gene expression, as determined by qPCR from whole islets (which contain ∼80% beta cells), was downregulated by 67% in KO *versus* littermate control islets after tamoxifen induction, while β-arrestin 1 levels were not significantly altered (Figure 1B). The weight, fasting and fed glycaemia of male or female mice on chow diet did not differ between genotypes (Supplemental Figure 1B, C). There were also no detectable differences in islet ultrastructure or insulin granule density as assessed by transmission electron microscopy (Supplemental Figure 1D), confirming previously reported data (25).

**Figure 1.**
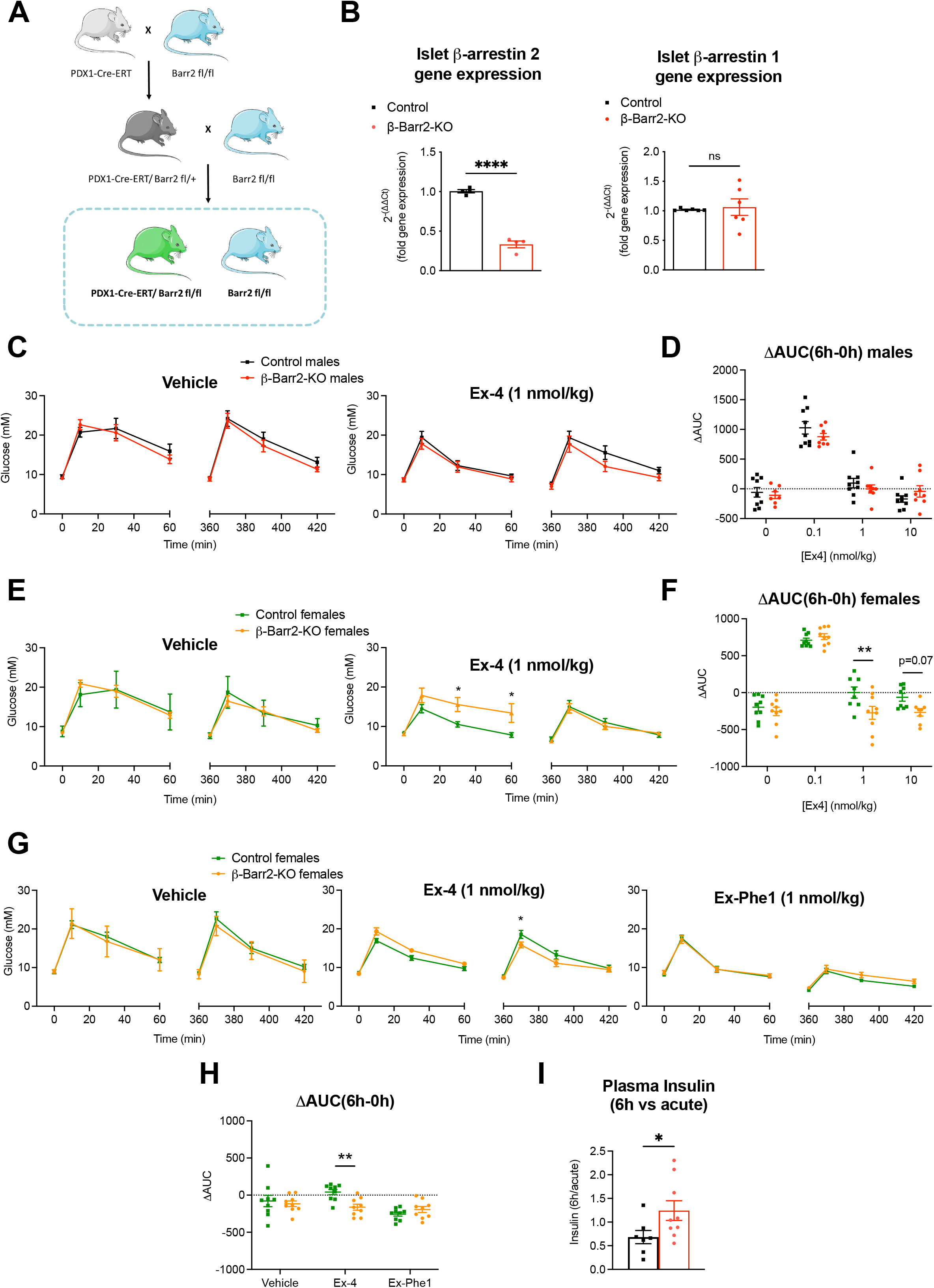
*In vivo* GLP-1R agonist responses in adult beta cell-selective β-arrestin 2 KO *vs* control mice on chow diet. IPGTTs (2 g/kg glucose i.p.) were performed concurrently with, or 6 h after, i.p. administration of agonists or vehicle (saline). (**A**) Schematic representation of the generation of Pdx1-Cre-ERT Barr2 fl/fl and control Barr2 fl/fl mice. (**B**) Relative gene expression of β-arrestin 2 (n = 4) and β-arrestin 1 (n = 6) in adult beta cell-specific β-arrestin 2 (β-Barr2) KO *vs* control mice. (**C**) Glucose curves for vehicle or exendin-4 (Ex-4) administration at 1 nmol/kg in lean, male mice (n = 8 / genotype, age: 12-16 weeks). (**D**) Corresponding ΔAUCs (6h-0h) generated from data in (C) and Supplemental Figure 1E. (**E**) Glucose curves for vehicle or Ex-4 administration at 1 nmol/kg in lean, female mice (n = 9 / genotype, age: 12-16 weeks). (**F**) Corresponding ΔAUCs (6h-0h) generated from data in (E) and Supplemental Figure 1F. (**G**, **H**) Glucose curves (**G**) and corresponding ΔAUCs (6h-0h) (**H**) for vehicle, 1 nmol/kg Ex-4, or 1 nmol/kg Exendin-Phe1 (Ex-Phe1) administration in lean, female mice (n = 9 / genotype, age: 14-18 weeks). (**I**) Plasma insulin fold changes (6 h over acute) from control *vs* KO mice calculated from data in Supplemental Figure 1G. Comparisons were made with unpaired t-tests or two-way ANOVA with Sidak’s *post hoc* tests. *p<0.05, **p<0.01, ****p<0.0001 *vs* control group. Data are presented as mean ± SEM.

Noting that the anti-hyperglycaemic effect of GLP-1R agonists (GLP-1RAs) biased away from β-arrestin recruitment becomes more prominent later into the dosing window (5), we performed intraperitoneal glucose tolerance tests (IPGTTs) both acutely and 6 hours after administration of exendin-4 at a range of doses (0.1, 1, and 10 nmol/kg). While we observed no significant effects at either time point for any of the doses in lean male mice (Figure 1C, D, Supplemental Figure 1E), KO female mice on chow diet displayed markedly worse acute glycaemic responses following 1 nmol/kg exendin-4 administration, with this difference subsequently lost at 6 hours post-treatment (Figure 1E, Supplemental Figure 1F), leading to a significantly improved ΔAUC (6 hours minus acute) factor when compared with control female mice (Figure 1F). Additional studies in these mice further confirmed the β-arrestin KO effect observed for exendin-4, but this was not present with the same dose of exendin-phe1, an exendin-4 derivative biased away from β-arrestin recruitment (5) and hence potentially less reliant on β-arrestin 2 engagement (Figure 1G, H). Concomitantly, plasma insulin levels measured 10 minutes into the IPGTT experiments were not changed between the two genotypes in the acute setting but were significantly raised for beta cell β-arrestin 2 KO *versus* control animals at 6 hours post-treatment when normalised to their corresponding acute levels (Figure 1I, Supplemental Figure 1G).

We also assessed the effect of the beta cell-selective β-arrestin 2 deletion on the related glucose-dependent insulinotropic polypeptide receptor (GIPR). To this effect, we tested the anti-hyperglycaemic effects of the stable GIPR agonist D-Ala^2^-GIP using similar methodology in lean animals. Similar trends for improved KO responses at 6 hours *versus* acutely were observed, although they did not quite reach statistical significance (Supplemental Figure 2A, B). Additionally, we were unable to replicate the exendin-4 results in a whole body, tamoxifen-inducible β-arrestin 2 KO model (R26-Cre-ERT2/Barr2 fl/fl), likely due to the compensatory effects of β-arrestin 2 deletion from different tissues, which, combined, might ablate the phenotype of the beta cell-selective model (Supplemental Figure 2E-H).

Next, to investigate the identified phenotype in a model of diet-induced metabolic stress and T2D, we administered high-fat high-sucrose (HFHS) diet to both control and beta cell β-arrestin 2 KO mice. Interestingly, KO males gained more weight compared with control littermates after prolonged HFHS diet exposure (Supplemental Figure 3A). Fasting glycaemia was elevated in HFHS-fed beta cell β-arrestin 2 KO males but not in females, with no observed changes in fed glycaemia between genotypes (Supplemental Figure 3B). Additionally, increased beta cell mass and average islet sizes were observed in pancreata from KO *versus* control HFHS-fed animals, with no changes in alpha cell mass and a trend for the alpha-to-beta cell mass ratio to be reduced (Supplemental Figure 3C, D). Study of expression levels of beta cell ‘enriched’ and ‘disallowed’ genes (26, 27) revealed that ‘enriched’ genes such as *MafA or Ins2* (with a tendency for *Kcjn11*) were significantly upregulated in KO animals, while expression of ‘disallowed’ genes (*Acot7*, *Slc16a1*, *Ldha*) was not altered (Supplemental Figure 3E).

HFHS-fed KO animals showed worsening glucose tolerance under vehicle conditions compared to control mice, particularly in males. However, the glucose-lowering effect of exendin-4 after glucose challenge was significantly more pronounced in KO *versus* control males 6 hours after agonist injection (Figure 2A), while, in females, there was an acute defect in exendin-4 responses from KO animals which, as in lean conditions, was overcome at 6 hours post-agonist exposure (Figure 2B). Both effects led to similar differences in control *versus* KO ΔAUCs (Figure 2C) and became highly significant when pooled following normalisation against the corresponding vehicle results (Figure 2D). To interrogate whether these effects were also present in clinically relevant agonists, we performed an analogous study in HFHS-fed animals with the long-acting GLP-1RA semaglutide and the dual GLP-1R/GIPR agonist tirzepatide [known to elicit reduced β-arrestin recruitment at the GLP-1R but not at its preferentially binding GIPR (28)], and observed a similar profile of improved responses at 72 hours *versus* 24 hours post-agonist injection in KO *versus* control animals (Figure 2E, F), indicating that lack of β-arrestin 2 is beneficial for prolongation of pharmacologically relevant GLP-1RA action *in vivo*. Additionally, and as for chow-fed animals, plasma insulin levels assessed 10 minutes into the IPGTTs from HFHS-fed animals showed no significant changes between the two genotypes in the acute settings, but there was a significant increase in KO *versus* control animals at 6 hours post-agonists exposure when normalised to their corresponding acute levels (Figure 2G, Supplemental Figure 3F).

**Figure 2.**
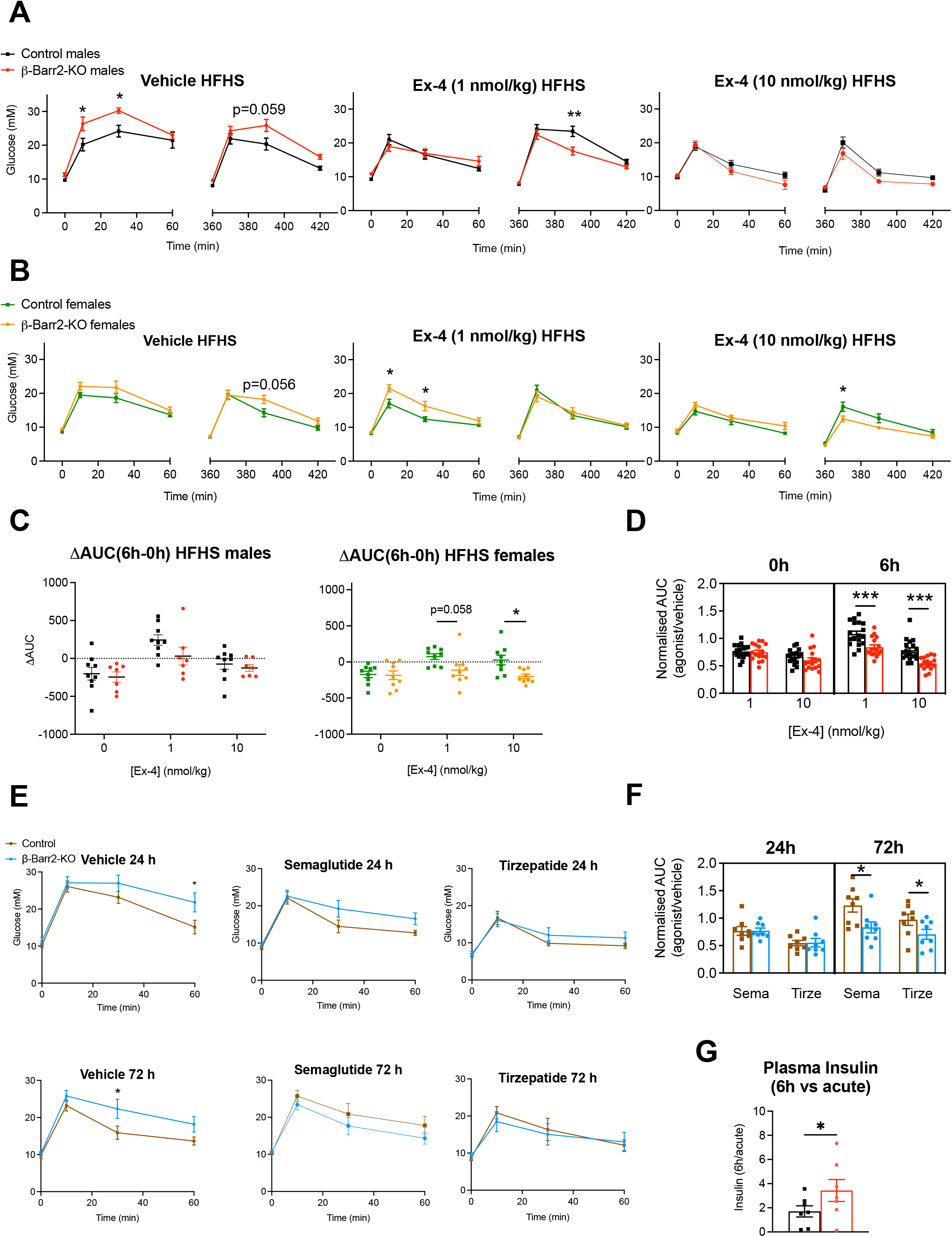
*In vivo* GLP-1R agonist responses in adult beta cell-selective β-arrestin 2 KO *vs* control mice on high-fat, high-sucrose (HFHS) diet. (**A**) Glucose curves for vehicle or Ex-4 administration at 1 and 10 nmol/kg in HFHS diet-fed male mice (n = 7-9 / genotype, duration of HFHS diet: 8-12 weeks, initiated at 8 weeks of age). (**B**) Glucose curves for vehicle or Ex-4 administration at 1 and 10 nmol/kg in HFHS diet-fed female mice (n = 9 / genotype, duration of HFHS diet: 8-12 weeks, initiated at 8 weeks of age). (**C**) ΔAUCs (6h-0h) for male and female HFHS diet-fed mice from (A, B). (**D**) Normalised AUCs demonstrating the effect of Ex-4 *vs* vehicle (AUC_Ex-4_/AUC_vehicle_) in HFHS diet-fed mice calculated from combined data presented in (A-C). (**E**) Glucose curves for vehicle, 10 nmol/kg semaglutide or 10 nmol/kg tirzepatide treatments; IPGTTs performed at 24 h or 72 h post-agonist i.p. injection in a mixed sex cohort (n = 8 / genotype, duration of HFHS diet: 8-12 weeks, initiated at 8 weeks of age). (**F**) Normalised AUCs demonstrating the agonist effect *vs* vehicle (AUC_agonist_/AUC_vehicle_) in HFHS diet-fed mice calculated from data presented in (E). (**G**) Plasma insulin fold changes (6 h over acute) from control *vs* KO mice calculated from data in Supplemental Figure 3F. Comparisons were made with unpaired t-tests, two- or three-way ANOVA with Sidak’s *post hoc* tests. *p<0.05, **p<0.01, ***p<0.001 *vs* control group. Data are presented as mean ± SEM.

### β-arrestin 2 regulates GLP-1R-triggered islet cAMP dynamics

For insight into the mechanism underpinning this contrasting acute *versus* prolonged glycaemic responses, we undertook further investigations in isolated islets. First, to investigate the role of β-arrestin 2 in the modulation of beta cell GLP-1R-dependent cAMP dynamics, we generated a beta cell-specific, tamoxifen-inducible mouse line conditionally expressing the CAMPER gene, encoding for the cAMP FRET biosensor ^T^Epac^VV^ (29) (Figure 3A). Time-lapse FRET microscopy experiments in control and beta cell-selective β-arrestin 2 KO CAMPER islets revealed significantly reduced acute cAMP responses to exendin-4 in KO *versus* control animals (Figure 3B). On the other hand, when islets were pre-treated with 1 nM exendin-4 overnight prior to washing and re-stimulation with GLP-1 to probe β-arrestin 2 relevance for GLP-1R desensitisation, we found that KO islets tended to produce higher cAMP responses compared with controls, therefore reversing the prior acute cAMP production defect (Figure 3C).

**Figure 3.**
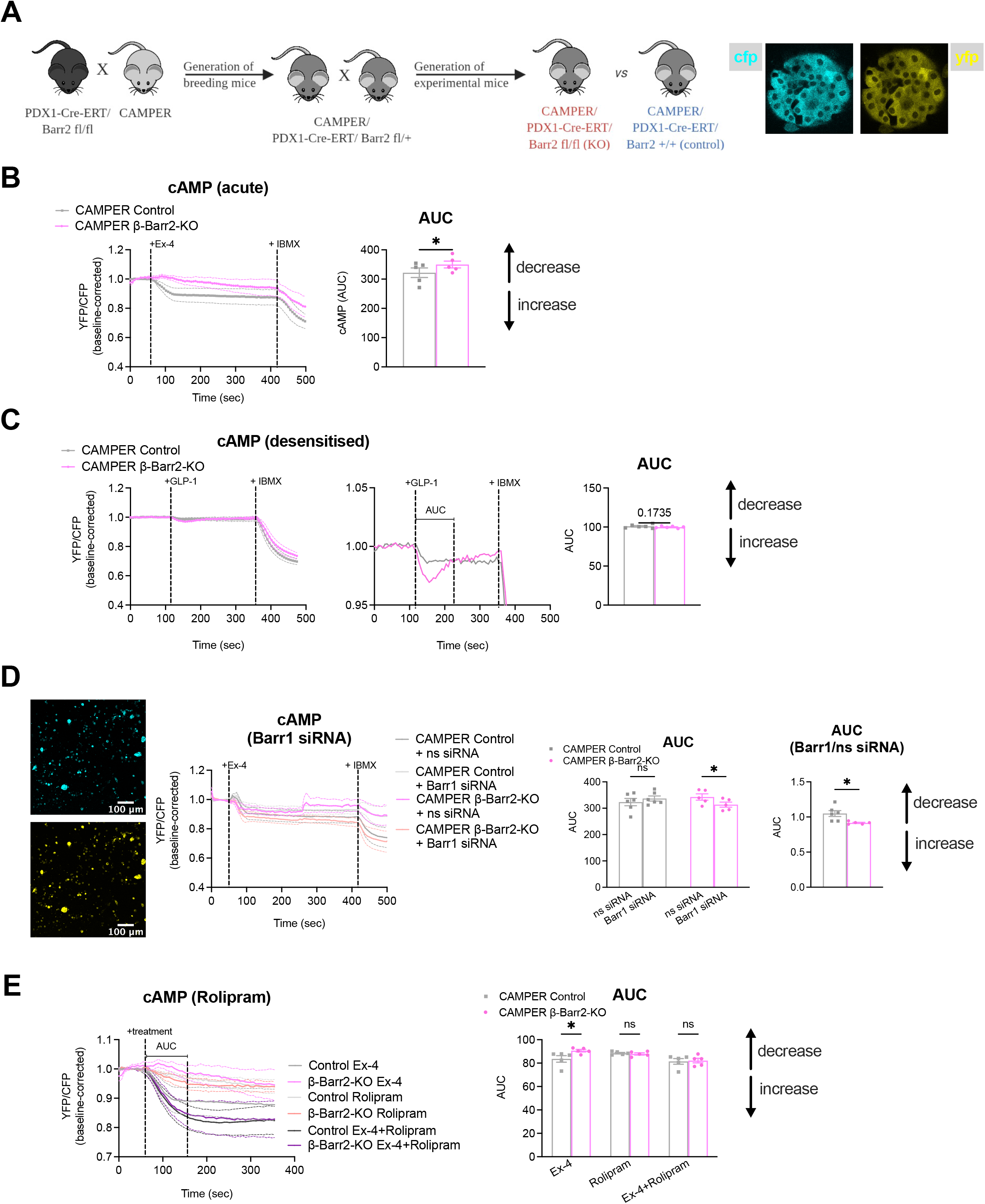
GLP-1R signalling alterations in adult beta cell-selective β-arrestin 2 KO *vs* control islets. (**A**) Schematic representation of mouse generation and representative images of the CFP and YFP channels from a CAMPER islet. (**B**) Changes in FRET (YFP/CFP) over time in isolated beta cell-selective β-arrestin 2 (β-Barr2) KO *vs* control CAMPER islets in response to 100 nM Ex-4 followed by 100 μM IBMX; AUCs calculated for the Ex-4 treatment period (n = 5 / genotype). (**C**) Changes in FRET (YFP/CFP) over time in isolated β-Barr2 KO *vs* control CAMPER islets pre-treated with 1 nM Ex-4 for 16 h (overnight) in response to 1 nM GLP-1 stimulation followed by 100 μM IBMX; AUCs calculated for the indicated GLP-1 treatment period (n = 6 / genotype). (**D**) Representative images of the CFP and YFP channel and changes in FRET (YFP/CFP) over time in cells from dispersed β-Barr2 KO *vs* control CAMPER islets pre-treated with non-specific (ns) control or β-arrestin 1 (Barr1) siRNA for 72 h in response to 100 nM Ex-4 followed by 100 μM IBMX; AUCs calculated for the Ex-4 treatment period (n = 5-6 / genotype). Corresponding Barr1 over control siRNA-treated islet fold changes included for each condition. (**E**) Changes in FRET (YFP/CFP) over time in isolated β-Barr2 KO *vs* control CAMPER islets in response to 100 nM Ex-4, 10 μM PDE4 inhibitor rolipram or a combination of the two; AUCs calculated for the indicated treatment period (n = 5 / genotype). Comparisons were made with t-tests, two-way ANOVA, or mixed-effects model with Sidak’s *post hoc* tests. *p<0.05 *vs* control group. Data are presented as mean ± SEM.

The cAMP phenotype was validated in our original non-CAMPER-expressing mouse model using an HTRF assay in response to a range of exendin-4 concentrations, where we again detected an acute cAMP defect in beta cell β-arrestin 2 KO conditions, with Emax and LogEC50 parameters reduced in both chow- and HFHS-fed mouse islets (Supplemental Figure 4A, B). We additionally performed time-lapse cAMP imaging experiments in islets infected with a baculovirus expressing the cAMP Difference Detector in situ (cADDis) biosensor (30), and, again in agreement with our CAMPER results, cADDis-transduced KO islets displayed a significant defect in acute cAMP *versus* control islets (Supplemental Figure 4C) but no difference in GLP-1 responses from desensitised islets (Supplemental Figure 4D).

We next hypothesised that, as the level of β-arrestin 1 is not reduced in the beta cell-selective β-arrestin 2 KO islets (Figure 1B), and GLP-1R has been previously shown to recruit β-arrestin 1 as well as 2 in beta cell lines (14), compensatory binding to this β-arrestin isoform in absence of the normally most abundant β-arrestin 2 might be responsible for the observed suboptimal GLP-1R cAMP production and/or accumulation. To investigate this, we tested the effect of RNAi-mediated β-arrestin 1 knockdown in dispersed (to allow good siRNA access) beta cell β-arrestin 2 KO and control CAMPER islets. Time-lapse FRET imaging experiments did indeed reveal that β-arrestin 1 knockdown restores normal acute cAMP responses to exendin-4 in KO CAMPER islets (Figure 3D), indicating that the previously identified acute cAMP defect is mediated by compensatory β-arrestin 1 action in the absence of β-arrestin 2. We reasoned that this could potentially result in changes in GLP-1R acute desensitisation, including, perhaps, differences in the control of cAMP degradation due to variations in the capacity for phosphodiesterase (PDE) recruitment to the receptor between the two β-arrestin isoforms (31). To test this hypothesis, and as PDE4 is the dominant isoform in beta cells (32, 33), we next evaluated the effect of rolipram, a specific PDE4 inhibitor (34), on the capacity for cAMP generation from beta cell-specific β-arrestin 2 KO and control CAMPER islets. Indeed, addition of rolipram normalised acute exendin-4-induced cAMP responses in KO islets (Figure 3E), implicating changes in PDE4 action in the acute cAMP defect under beta cell β-arrestin 2 KO conditions.

### Beta cell β-arrestin 2 deletion modifies GLP-1RA-induced intra-islet Ca^2+^ dynamics and connectivity

Changes in intracellular free Ca^2+^ have been described as another downstream signalling readout of GLP-1R activation, controlled mainly by G_αs_ but also by G_αq_ protein coupling (35). Intracellular calcium rises are more distal within the signalling pathway and linked to insulin granule exocytosis. We used the calcium dye Cal-520 AM and time-lapse fluorescence microscopy to investigate intracellular calcium dynamics in response to acute exendin-4 stimulation in beta cell β-arrestin 2 KO *versus* control islets from either chow- or HFHS-fed animals. On chow diet, both islet types displayed similar exendin-4-induced calcium rises at 6 mM glucose, although response to a subsequent 11 mM glucose challenge was blunted in KO islets (Figure 4A). However, in keeping with the *in vivo* observation that HFHS feeding accentuates β-arrestin 2 effects on exendin-4 responses, islets isolated from HFHS-fed KO animals displayed significantly blunted responses to exendin-4 throughout the acquisition compared to controls (Figure 4B).

**Figure 4.**
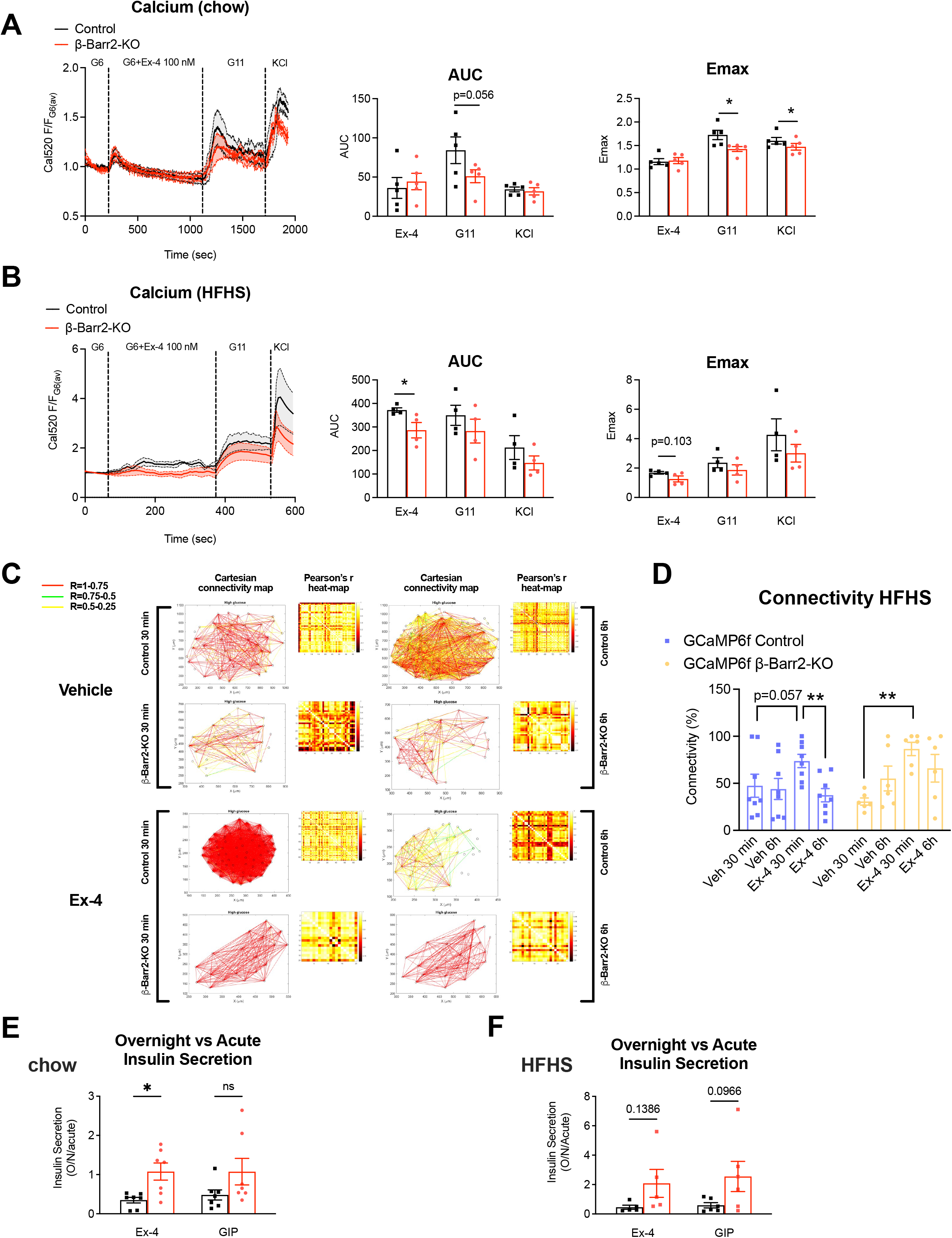
Downstream effects of GLP-1R signalling alterations in adult beta cell-selective β-arrestin 2 KO *vs* control islets. (**A**) Calcium traces and corresponding AUCs and Emax for the indicated treatments in beta cell-selective β-arrestin 2 (β-Barr2) KO *vs* control islets isolated from animals on chow diet (n = 5 / genotype). (**B**) Calcium traces and corresponding AUCs and Emax for indicated treatments in β-Barr2 KO *vs* control islets isolated from animals on HFHS diet (n = 4 / genotype). (**C**, **D**) Representative cartesian connectivity maps (C) and percentage of connectivity (D) for β-Barr2 KO *vs* control GCaMP6f islets from HFHS diet donors implanted in the anterior chamber of the eye of WT acceptors on HFHS diet, which received 2 g/kg glucose i.p. with Ex-4 10 nmol/kg or saline (vehicle) (n = 6-8 islets / genotype). (**E**, **F**) Insulin secretion fold changes [overnight over acute (1h)] from control vs β-Barr2 KO islets on chow (E) or HFHS diet (F) calculated from data in Supplemental Figure 4I, J for the indicated agonist. Comparisons between two experimental groups were made with t-tests, one- or two-way ANOVA or mixed-effects model with Sidak’s *post hoc* tests. *p<0.05, **p<0.01. Data are presented as mean ± SEM.

Next, to investigate calcium responses *in vivo*, a mouse model of the genetically encoded calcium indicator GCaMP6f (36) was crossed with β-arrestin 2 KO mice to generate beta cell-specific GCaMP6f β-arrestin 2 KO and wild type (WT) control mice, and islets from these mice isolated and implanted in the anterior chamber of the eye of WT acceptor mice. This platform supported longitudinal *in vivo* imaging of calcium responses as previously shown (37, 38) using mildly elevated *in vivo* glucose levels. Experiments were carried out under both chow and HFHS diet conditions, with islet donors and acceptors matched for diet type. Islet calcium ‘waves’ in response to treatments were classified into 4 activity categories, with 1 representing the least active and 4 the most active category. In agreement with previous results from diabetic islets (37), implanted islets from HFHS-fed mice displayed lower activity compared with their chow diet counterparts (Supplemental Figure 4E). The number of islets examined did not permit for statistical tests, but there was a trend for higher activity indices in HFHS islets from beta cell β-arrestin 2 KO *versus* control mice. Wave characteristics, including wavelength, amplitude, and full width at half-maximum (FWHM) did not differ between treatments and genotypes for chow diet islets (Supplemental Figure 4F). For HFHS diet, the amplitude of calcium waves was significantly increased in KO *versus* control islets at the 6-hour but not at the 30-minute time-point for both vehicle and 10 nmol/kg exendin-4 administration, while the wavelength and FWHM were not changed (Supplemental Figure 4G). The percentage of connectivity of single cells within islets, a measure of coordinated intra-islet responses, was not significantly different between genotypes after exendin-4 administration on chow diet (Supplemental Figure 4H).

However, under HFHS diet conditions, connectivity initially increased under both genotypes following acute exendin-4 stimulation, then waned over time from 30 minutes to 6 hours post-exendin-4 injection in control islets, but, conversely, a high percentage of connectivity was retained at 6 hours post-exendin-4 administration in KO islets (Figure 4C, D), suggesting that loss of coordinated calcium responses after sustained exendin-4 exposure is overcome *in vivo* by deletion of β-arrestin 2.

We next performed *ex vivo* insulin secretion assays from isolated islets, with 1-hour exendin-4 incubations used as acute, and overnight (16-hour, cumulative) incubations as prolonged readouts. Here we observed reduced acute exendin-4-induced insulin secretion (*versus* 11 mM glucose) for beta cell β-arrestin 2 KO islets from animals on either diet type *versus* controls. However, this was reversed overnight, with this temporal trajectory conveniently expressed as “overnight over acute”, an effect significant for islets from mice on chow diet and close to significance for HFHS diet (Figure 4E, F, Supplemental Figure 4I, J). Parallel experiments, depicted in the same figure, were performed using the GIPR agonist GIP, where again we observed the same trend, which, in agreement with our *in vivo* results with GIP-D-Ala^2^, did not quite reach statistical significance, suggesting a lesser dependence of GIPR on β-arrestin 2.

### Islet GLP-1R trafficking is perturbed following β-arrestin 2 deletion

We next assessed the potential contribution from alterations in GLP-1R trafficking to the prolongation of exendin-4-induced signalling in the absence of β-arrestin 2. First, to control for potential differences due to changes in basal cell surface GLP-1R expression, we quantified surface GLP-1R levels from control and beta cell β-arrestin 2 KO islets by labelling these with the fluorescent antagonist exendin-9-TMR, with no differences observed in surface GLP-1R levels between the two genotypes (Figure 5A). We next determined endogenous GLP-1R trafficking profiles in beta cell β-arrestin 2 KO *versus* control islets using the previously characterised fluorescent agonist exendin-4-TMR (39) as a proxy for GLP-1R localisation. We quantified GLP-1R internalisation in response to 1hour exendin-4-TMR exposure by measuring TMR fluorescence levels, using acetic acid buffer wash treatment to strip any remaining exendin-4-TMR bound to non-internalised/cell surface receptors, both for chow- and HFHS islets (Figure 5B, Supplemental Figure 5A). Of note, the acid wash did not significantly reduce TMR signal, indicating that exendin-4-TMR is predominantly internalised under these conditions. We also noted that islets from HFHS-fed mice had generally lower exendin-4-TMR binding capacity than chow islets (Supplemental Figure 5A). We were unable to detect any differences in GLP-1R internalisation between KO and control islets, suggesting that, as previously observed in cell lines (13), β-arrestin 2 does not play a significant role in GLP-1R endocytosis. Next, we assessed the propensity for GLP-1R plasma membrane recycling by incubating control or KO islets with exendin-4-TMR for 3 hours after a prior 1-hour incubation with unlabelled exendin-4 to trigger maximal receptor internalisation in both chow- and HFHS-fed mouse islets. In this assay, re-emergence of agonist-internalised GLP-1R at the cell surface is detected by subsequent binding and re-uptake of exendin-4-TMR. While no differences in GLP-1R recycling were detected at the 3-hour time-point for chow islets, HFHS KO islets displayed a small but significant reduction in GLP-1R recycling following exendin-4 stimulation (Figure 5C, Supplemental Figure 5B). We repeated the recycling assay in chow islets for 6 hours of receptor recycling to assess any possible changes that might only become apparent within longer periods (Figure 5D, Supplemental Figure 5C), and, under these conditions, a small but significant reduction in GLP-1R recycling became indeed apparent in KO *versus* control islets from chow-fed mice.

**Figure 5.**
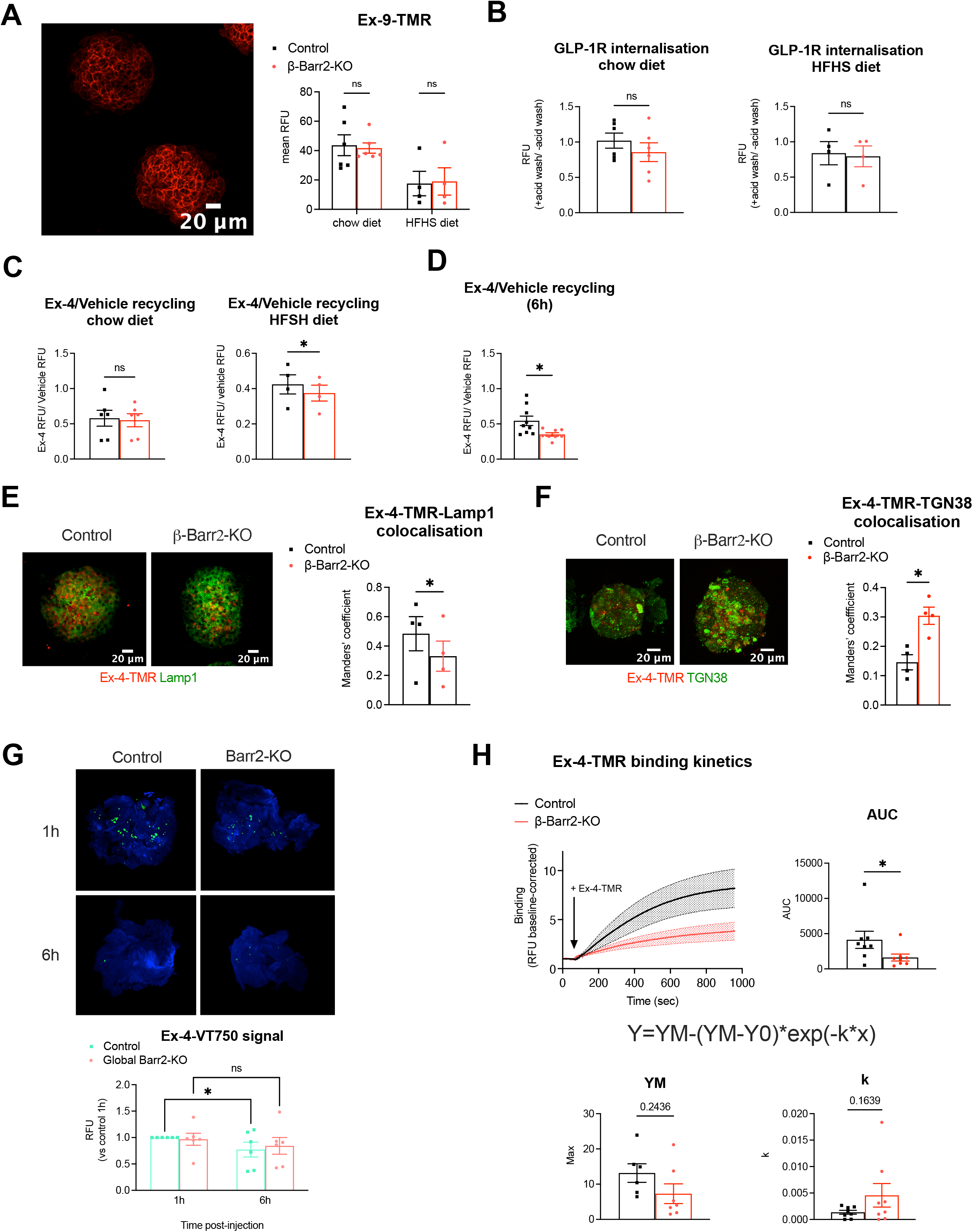
GLP-1R trafficking alterations in adult beta cell-selective β-arrestin 2 KO *vs* control islets. (**A**) Representative image of maximum intensity projections from confocal z-stacks, and quantification of TMR fluorescence in beta cell-selective β-arrestin 2 (β-Barr2) KO *vs* control islets treated with 100 nM Ex-9-TMR for 30 min as a measure of surface GLP-1R levels in mice on chow diet (n = 6 / genotype) or HFHS diet (n = 4 / genotype). (**B**) Ex-4-TMR fluorescence at 1 h (depicted as fold with *vs* without acid wash), used as a surrogate measure for GLP-1R internalisation in β-Barr2 KO *vs* control islets from chow- and HFHS-fed mice, calculated from data in Supplemental Figure 5A. (**C**) Normalised Ex-4 over vehicle GLP-1R recycling at 3 h in β-Barr2 KO *vs* control islets from chow- and HFHS-fed mice, calculated from data in Supplemental Figure 5B. (**D**) Normalised Ex-4 over vehicle GLP-1R recycling at 6 h in β-Barr2 KO *vs* control islets from chow-fed mice, calculated from data in Supplemental Figure 5C. (**E**) Representative images of maximum intensity projections from confocal z-stacks of β-Barr2 KO *vs* control islets labelled for Lamp1 (green) after treatment with 100 nM Ex-4-TMR (red) for 3 h. Manders’ coefficient of co-localisation for Ex-4-TMR with Lamp1 signal is shown (n = 4 / genotype). (**F**) Representative images of maximum intensity projections from confocal z-stacks of β-Barr2 KO *vs* control islets labelled for TGN38 (green) after treatment with 100 nM Ex-4-TMR (red) for 3 h. Manders’ coefficient of co-localisation for Ex-4-TMR with TGN38 signal is shown (n = 4 / genotype). (**G**) Representative images from optical projection tomography data from pancreata extracted from whole-body β-arrestin 2 (Barr2) KO and control animals injected i.p. with 100 nmol/kg Ex-4-VT750 1 h or 6 h post-injection. Mean Ex-4-VT750 islet fluorescence depicted for each condition (n = 6 / genotype). (**H**) GLP-1R binding affinity to 100 nM Ex-4-TMR in β-Barr2 KO *vs* control islets pre-treated with metabolic inhibitors (n = 8 / genotype). AUC, maximum value (YM), and rate constant (k) from fitted curves are depicted for each genotype. Comparisons were made using t-tests, or two-way ANOVA with Sidak’s *post hoc* tests. *p<0.05 *vs* control group. Data are presented as mean ± SEM.

We next assessed the co-localisation between exendin-4-TMR and LAMP1, a lysosomal marker, as well as TGN38, a marker for the trans-Golgi network (TGN), in islets after 3 hours of exendin-4-TMR stimulation (Figure 5E, F). Here we found significantly reduced co-localisation between fluorescent exendin-4 and LAMP1, as well as increased exendin-4-TMR co-localisation with TGN38 in KO *versus* control islets, suggesting reduced capacity for GLP-1R trafficking to lysosomal compartments and increased propensity for redirection to the TGN in the absence of β-arrestin 2.

Next, to assess *in vivo* GLP-1R agonist accumulation within the pancreas following exendin-4 exposure, we performed 3D optical projection tomography imaging and signal quantification from optically cleared whole pancreata extracted from global β-arrestin 2 KO or control mice previously injected with the near infrared exendin-4 derivative exendin-4-VivoTag 750 for either 1 hour or 6 hours prior to intracardial fixation and pancreas extraction (Figure 5G). Results showed a significant loss of signal from control mice pancreata at the 6-hour over the 1-hour period, while signal was instead maintained in β-arrestin 2 KO samples.

Finally, we evaluated the GLP-1R capacity for binding to exendin-4-TMR in islets from beta cell-specific β-arrestin 2 KO *versus* control mice imaged by time-lapse confocal microscopy in the presence of metabolic inhibitors to inhibit receptor endocytosis (Figure 5H). Fitted curve results showed that while kinetic parameters were not altered, there was a significant reduction in AUCs in beta cell β-arrestin 2 KO compared with control islets, indicating the existence of a defect in GLP-1R exendin-4-TMR binding capacity for beta cells lacking β-arrestin 2, a phenotype that might contribute to the observed deficit in acute GLP-1R signalling from these islets.

### Impact of β-arrestin 2 deletion on GLP-1R trafficking and signalling in a beta cell model

After characterisation of GLP-1R responses in adult beta cell β-arrestin 2 KO mice and primary islets, we next generated an *in vitro* beta cell model for a more detailed examination of molecular mechanisms associated with the changes observed in GLP-1R trafficking and signalling. We first verified plasma membrane recruitment of β-arrestin 2 following GLP-1R stimulation in our chosen model, namely INS-1 832/3 rat insulinoma cells (Supplemental Figure 6A, B). Time-lapse spinning disk microscopy experiments in cells stably expressing SNAP-tagged human GLP-1R and transiently transfected with a β-arrestin 2-GFP construct demonstrated that β-arrestin 2 is indeed recruited from the cytoplasm to the plasma membrane, where it co-localises with SNAP-GLP-1R within 5 minutes of exendin-4 stimulation.

Next, we generated a lentiviral CRISPR/Cas9-derived INS-1 832/3 cell subline in which the β-arrestin 2 gene was ablated. The resulting β-arrestin 2 knockdown (KD) cells displayed a 51% reduction in β-arrestin 2 expression compared with control cells generated in parallel with a non-targeting construct (Figure 6A). Using this cell model, we assessed the degree of GLP-1R plasma membrane *versus* endosomal signalling by implementing a NanoBiT assay based on the Nb37 (40), a nanobody which specifically binds to active G_αs_ proteins following GPCR stimulation (41). In agreement with our previously detected acute cAMP defect in beta cell β-arrestin 2 KO islets, we measured a small but significant reduction in Emax, with no change in logEC50, for exendin-4-induced GLP-1R plasma membrane signalling in β-arrestin 2 KD compared with control cells, with no changes associated with the degree of endosomal signalling (Figure 6B, C). A similar experiment performed with the GIPR agonist D-Ala^2^-GIP in the same cells did not reveal any significant changes in GIPR plasma membrane or endosomal signalling in β-arrestin 2 KD *versus* control cells (Supplemental Figure 7A, B).

**Figure 6.**
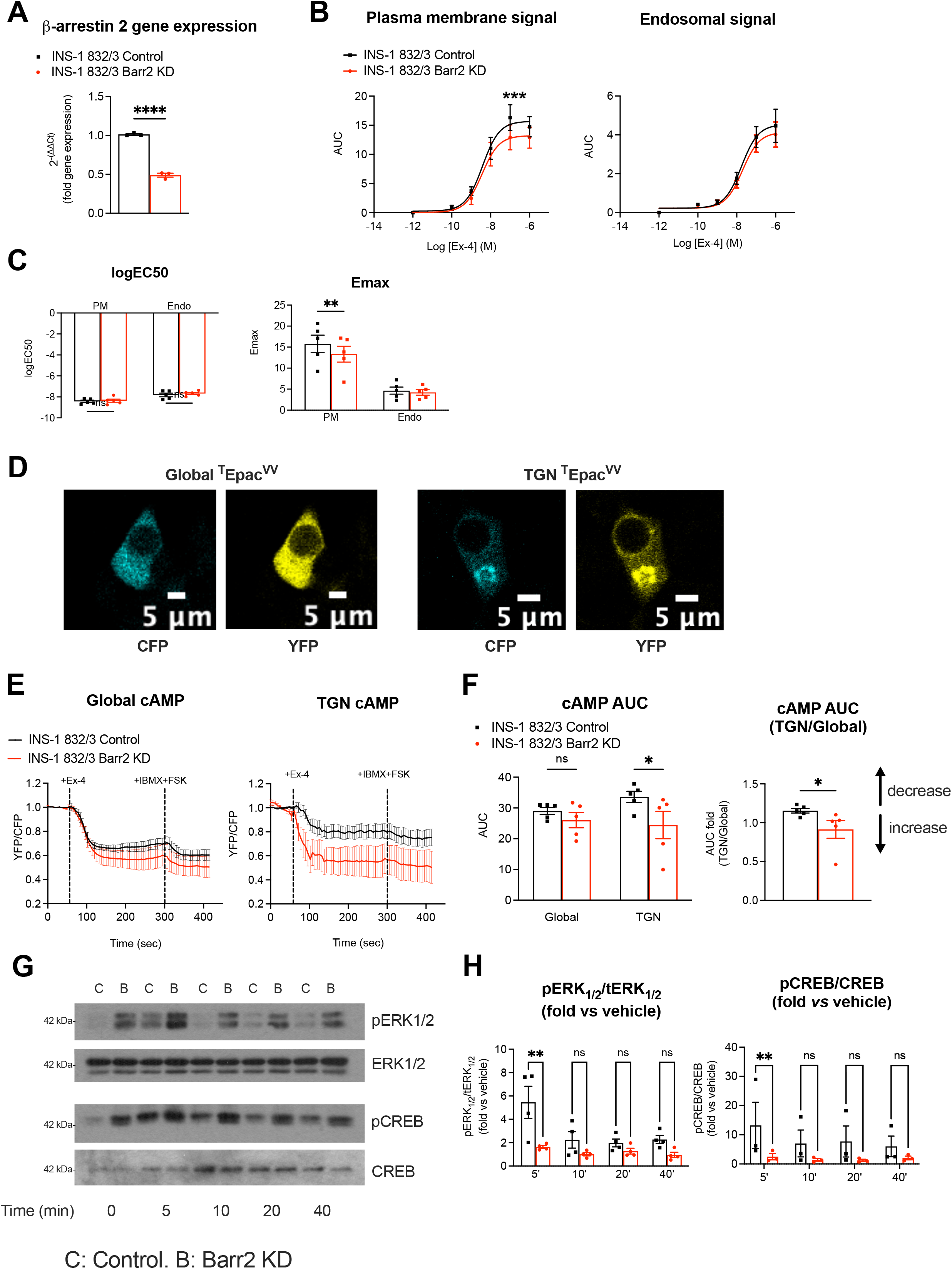
Spatiotemporal signalling profiles of INS-1 832/3 β-arrestin 2 KD *vs* control beta cells. (**A**) Relative β-arrestin 2 gene expression in β-arrestin 2 (Barr2) KD *vs* control INS-1 832/3 cells (n = 3). (**B**) Ex-4 dose-response AUC curves for Nb37-SmBiT and CAAX-LgBiT (plasma membrane) or Endofin-LgBiT (endosomal) signal complementation assays (n = 5). (**C**) LogEC50 and Emax values calculated from (B). (**D**) Representative images of CFP and YFP channels from INS-1 832/3 cells transfected either with global (cytoplasmic) or TGN-targeted ^T^Epac^VV^ cAMP biosensor constructs. (**E**) Changes in FRET (YFP/CFP) over time in INS-1 832/3 Barr2 KD *vs* control cells expressing global or TGN-targeted ^T^Epac^VV^ cAMP biosensor constructs in response to 100 nM Ex-4 followed by 100 μM IBMX + 10 μM forskolin (n = 5). (**F**) AUCs calculated for the Ex-4 treatment period from (E). TGN-localised normalised to global cAMP AUCs are depicted for each cell type. (**G**) Representative blots for pERK1/2, total ERK1/2, pCREB, and total CREB in Barr2 KD and control INS-1 832/3 cells after treatment with 100 nM Ex-4 for the indicated time-points. (**H**) Quantification of pERK1/2 over total ERK1/2, and pCREB over total CREB (fold *vs* vehicle) using densitometry analysis (n = 4 for pERK1/2 and n = 3 for pCREB). Data from Supplemental Figure 7C normalised to vehicle levels for each cell type. Comparisons were made using t-tests, or two-way ANOVA with Sidak’s *post hoc* tests. *p<0.05, **p<0.01, ***p<0.001, ****p<0.0001 *vs* control group. Data are presented as mean ± SEM.

Following our *ex vivo* trafficking results showing reduced GLP-1R targeting to lysosomal compartments but increased TGN localisation in β-arrestin 2 KO islets, we next assessed the level of signalling specifically from this latter compartment with time-lapse FRET microscopy using a ^T^Epac^VV^-based cAMP biosensor modified *in house* to localise specifically to the TGN (Figure 6D). Quantification of cAMP production after exendin-4 stimulation with both global (cytoplasmic) and TGN-targeted ^T^Epac^VV^ biosensors demonstrated a significant increase in TGN over global cAMP in β-arrestin 2 KD *versus* control cells (Figure 6E, F). Downstream signal transmission was next assessed by Western blot analysis of ERK1/2 and CREB phosphorylation in both control and β-arrestin 2 KD cells at different times of exendin-4 exposure. This experiment showed increased basal phospho-ERK1/2 (and a similar trend for basal phospho-CREB) but reduced ERK1/2 and CREB phosphorylation fold increases at 5 minutes post-exendin-4 stimulation in β-arrestin 2 KD *versus* control cells (Figure 6G, H, Supplemental Figure 7C).

We next evaluated the endogenous GLP-1R cell surface levels in both cell lines by labelling them with exendin-9-TMR and, as for primary islets, we found no differences in surface GLP-1R between control and β-arrestin 2 KD cells (Supplemental Figure 7D). We then performed a comprehensive assessment of GLP-1R trafficking in these cells by using NanoBRET subcellular localisation assays based on expression of specific Rab GTPase bystanders at different intracellular locations (Supplemental Figure 7E), revealing no differences in GLP-1R plasma membrane or Rab5-positive early endosome localisation in response to increasing concentrations of exendin-4 between both cell types, but, in agreement with our primary islet recycling results, a clear decrease in GLP-1R localisation to Rab11-positive recycling endosomes, associated with reduced Emax values in β-arrestin 2 KD compared with control cells (Figure 7A, B).

**Figure 7.**
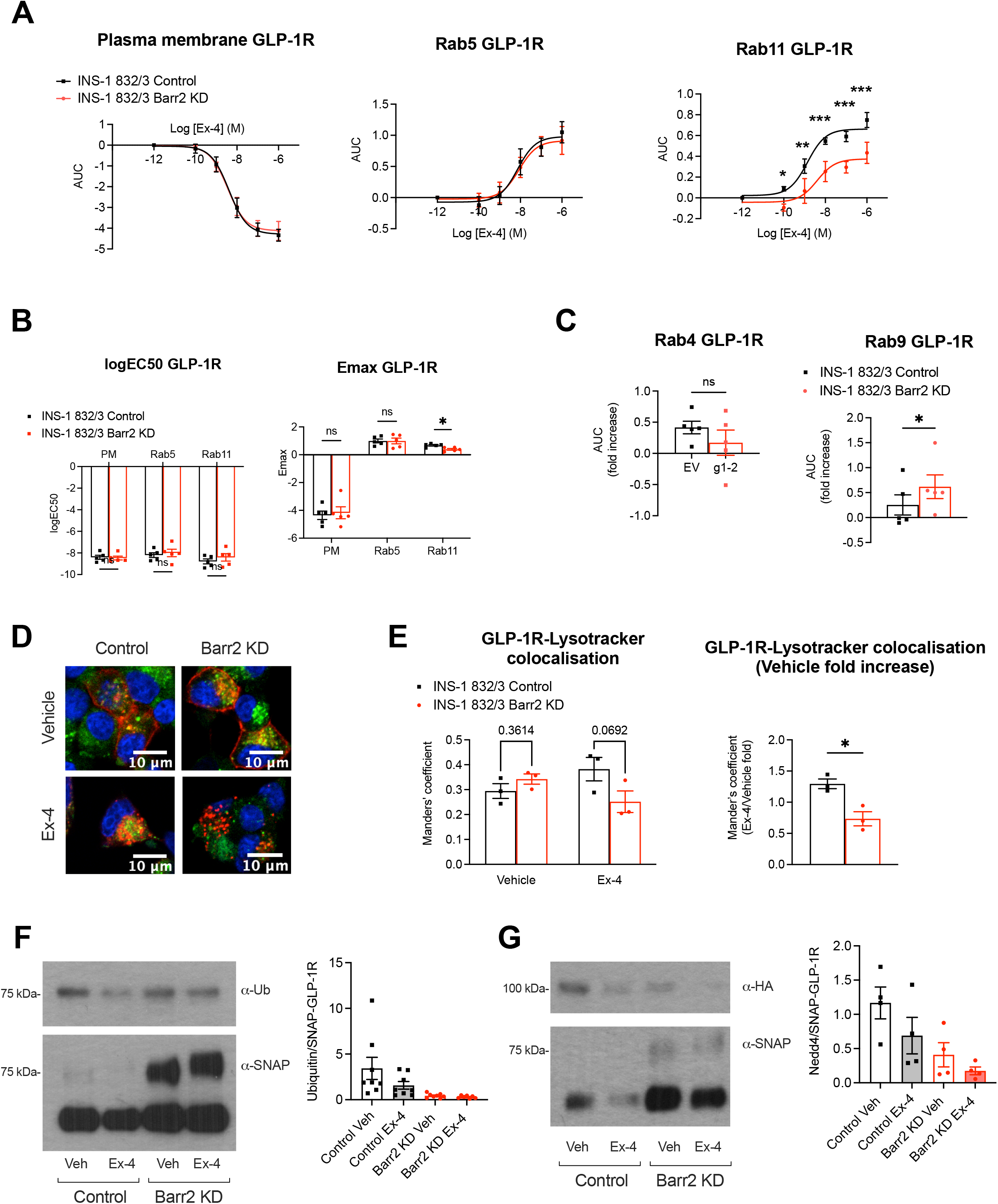
Associated changes in GLP-1R trafficking and ubiquitination in INS-1 832/3 β-arrestin 2 KD *vs* control beta cells. (**A**) Ex-4 dose-response AUC curves for SNAP-GLP-1R-NLuc and KRAS-Venus (plasma membrane), Rab5-Venus (early endosomes), and Rab11-Venus (recycling endosomes) NanoBRET assays (n = 5). (**B**) LogEC50 and Emax values calculated from (A). (**C**) AUC for NanoBRET responses to 100 nM Ex-4 using SNAP-GLP-1R-NLuc and Rab4-Venus (fast recycling endosomes) or Rab9-Venus (endosome-to-TGN). (**D**) Representative images of SNAP-GLP-1R-expressing Barr2 KD and control cells labelled with SNAP-Surface 647 (red) and treated with 100 nM Ex-4 or vehicle for 3 h and 1 μM LysoTracker Red DND-99 (green) for the last 5 min of the incubation period to label the lysosomes; Nuclei, DAPI (blue). (**E**) Manders’ coefficient of co-localisation for SNAP-GLP-1R with LysoTracker Red DND-99 in INS-1 832/3 Barr2 KD *vs* control cells (n = 3). Ex-4 over vehicle fold values depicted as a surrogate measure of agonist-induced SNAP-GLP-1R degradation (n = 3). (**F**) Representative blots showing SNAP-GLP-1R and corresponding ubiquitination levels following SNAP-GLP-1R immunoprecipitation from INS-1 832/3 SNAP-GLP-1R Barr2 KD *vs* control cells under vehicle conditions of after stimulation with 100 nM Ex-4 for 10 min, with quantification of ubiquitin over SNAP levels for the different conditions shown (n = 8). (**G**) Representative blots showing SNAP-GLP-1R and HA-NEDD4 levels following SNAP-GLP-1R immunoprecipitation from INS-1 832/3 SNAP-GLP-1R Barr2 KD *vs* control cells under vehicle conditions of after stimulation with 100 nM Ex-4 for 10 min, with quantification of HA over SNAP levels for the different conditions shown (n = 4). Comparisons were performed using t-tests, or one- or two-way ANOVA with Sidak’s *post hoc* tests. *p<0.05, **p<0.01, ***p<0.001. Data are presented as mean ± SEM.

We also investigated any differences associated with GLP-1R localisation to either Rab4- or Rab9-positive compartments, highlighting a fast recycling route or the retrograde endosome-to-Golgi transport, respectively, and while we found no effect on GLP-1R localisation to Rab4 endosomes, the measured significantly increased GLP-1R recruitment to Rab9-positive compartments in β-arrestin 2 KD *versus* control cells (Figure 7C), suggesting that these cells recapitulate the trafficking phenotype obtained in beta cell β-arrestin 2 KO primary islets, with increased GLP-1R re-routing to the TGN. We also performed GLP-1R lysosomal localisation studies in control and β-arrestin 2 KD cells transiently expressing SNAP-GLP-1R labelled with a fluorescent SNAP-Surface probe and Lysotracker under both vehicle and exendin-4-stimulated conditions (Figure 7D, E), and, as for islets, found significantly reduced lysosomal targeting of exendin-4-stimulated GLP-1Rs in β-arrestin 2 KD *versus* control cells.

Finally, we investigated the degree of GLP-1R ubiquitination as a potential mechanism that might explain the post-endocytic trafficking differences observed between β-arrestin 2 KD and control cells. To this effect, we performed GLP-1R immunoprecipitation experiments in INS-1 832/3 control and β-arrestin 2 KD cells modified to stably express SNAP/FLAG-tagged human GLP-1Rs (Figure 7F, Supplemental Figure 7F). Results in control cells unexpectedly showed that the GLP-1R is constitutively ubiquitinated under vehicle conditions and subsequently undergoes partial deubiquitination following exendin-4 stimulation. They also revealed a pronounced reduction in the level of GLP-1R ubiquitination in β-arrestin 2 KD cells, present under both vehicle and stimulated conditions, so that further de-ubiquitination in response to exendin-4 is no longer significant. We have previously identified, in an RNAi screen for factors involved in regulating exendin-4-stimulated insulin secretion (42), the E3 ubiquitin ligase NEDD4 (neural precursor cell expressed, developmentally downregulated-4), best known to mediate β-arrestin 2-dependent ubiquitination and endo-lysosomal sorting of GPCRs following ligand stimulation (43, 44), as a factor involved in the maintenance of exendin-4-stimulated insulin secretion. More recently, we have again identified NEDD4 as a GLP-1R interactor in a mass spectrometry analysis of the receptor interactome in beta cells (unpublished results). Therefore, we analysed whether the changes in GLP-1R ubiquitination in cells without β-arrestin 2 could be attributed to altered recruitment of this ubiquitin ligase to the receptor. To test this, we co-immunoprecipitated NEDD4 with GLP-1R from control and β-arrestin 2 KD cells stably expressing SNAP/FLAG-GLP-1R and transiently transfected with an HA-tagged NEDD4 construct to quantify the level of HA-NEDD4-SNAP/FLAG-GLP-1R association both in vehicle and exendin-4-stimulated conditions (Figure 7G, Supplemental Figure 7G). Results for NEDD4-GLP-1R interaction closely correlated with those for GLP-1R ubiquitination, as GLP-1R was found constitutively associated with NEDD4, with GLP-1R-NEDD4 interaction partially lost following exendin-4 stimulation. Furthermore, the level of GLP-1R-associated NEDD4 was significantly reduced in β-arrestin 2 KD *versus* control cells, a phenotype that, as for ubiquitination, was already present under vehicle conditions.

## Discussion

This study builds upon our previous cell line data suggesting GLP-1R signal prolongation following β-arrestin downregulation (17), although these experiments were performed after knockdown of both β-arrestin isoforms and were therefore not specific for β-arrestin 2. To date, there was very limited data on primary islets and none *in vivo* on the precise role of this important signalling regulator in modulating GLP-1R responses, despite the known capacity of biased GLP-1RAs with differential G protein over β-arrestin recruitment propensities to modify beta cell behaviours (3, 5). Although a previous report had suggested that islets from adult beta cell β-arrestin 2 deleted mice had normal *ex vivo* GLP-1R responses, this data was from acute experiments in islets extracted from lean male mice (25), a condition for which, in consonance, we also failed to detect any significant *in vivo* effects. However, after performing careful assessments, we were able to establish a clear *in vivo* phenotype in lean female mice, as well as in male and female mice fed a HFHS diet over 16 weeks, unveiling an acute signalling defect that progresses towards improved exendin-4 responses over longer stimulation periods in beta cell β-arrestin 2-deleted animals. While the reasons behind the observed sex differences on β-arrestin 2 dependency for GLP-1R responses have not been analysed here in detail, it is of note that males, which are well known to display (as also observed here) increased susceptibility to impaired glucose metabolism and T2D (45), are less reliant on β-arrestin 2 for their pharmacological GLP-1R responses, opening the door to potential sex differences in responses to incretin treatment, as previously detected for glucose-stimulated GLP-1 secretion responses (46).

The validity of our *in vivo* data is reinforced by our observations of concomitant increases in sustained over acute plasma insulin levels from β-arrestin 2 KO *versus* control mice, as well as by the lack of effect with exendin-phe1, an exendin-4-derived GLP-1RA with a single amino acid substitution biased away from β-arrestin recruitment (5), a feature that likely allows it to bypass both the positive and the negative effects of β-arrestin 2 on GLP-1R signalling. Importantly, the clinical relevance of our observations is clearly established as the currently leading, as well as the most promising pharmacological GLP-1RAs, namely semaglutide and tirzepatide (47), were both similarly affected to exendin-4 by the lack of beta cell β-arrestin 2. Interestingly, however, we could not replicate the *in vivo* phenotype in a whole-animal β-arrestin 2 KO model, suggesting that β-arrestin 2 modulation of GLP-1R action in beta cells is compensated by concomitant changes in the response of this or of other GPCRs within other tissues, resulting in a zero net effect in glucose handling. Additionally, while the effect of knocking out beta cell β-arrestin 2 on physiological islet responses has not been extensively characterised here, we have found, in agreement with previously published data (25), a significant glucoregulatory defect in HFHS-fed mice under vehicle conditions, suggesting a negative impact of absence of β-arrestin 2 on general beta cell function, which, in our study, was accompanied by increases in beta cell mass and average islet sizes, as well as by increased expression of beta cell ‘enriched’ genes, indicating lack of receptor signalling restraint and greater beta cell hypertrophy under metabolic stress in cells lacking β-arrestin 2.

We have additionally performed a comprehensive *ex vivo* study of the consequences of deleting β-arrestin 2 from primary beta cells on GLP-1R downstream signalling responses in islets, revealing an acute cAMP defect by three separate methods, subsequently overcome when probing for receptor desensitisation after sustained agonist exposure. Moreover, we have determined the involvement of β-arrestin 1 and the cAMP phosphodiesterase PDE4 on the establishment of this acute defect, as cAMP production was restored in islets following siRNA-mediated β-arrestin 1 knockdown or treated with the PDE4-specific inhibitor rolipram. These two effects are likely functionally linked, as β-arrestins can desensitize GPCRs not solely by homologous desensitisation, but also by recruiting specific PDE isoforms to control the rate of local cAMP degradation (31, 48, 49), so that abnormally-increased GLP-1R - β-arrestin 1 recruitment in the absence of β-arrestin 2 might result in a higher degree of cAMP dampening via augmented recruitment of PDE4. Additionally, we observed a reduction in acute GLP-1R binding to exendin-4-TMR in islets lacking beta cell β-arrestin 2: this could again represent a difference in behaviour between both arrestins, as β-arrestin 2 is known to induce higher agonist-binding affinity receptor conformations than β-arrestin 1 for certain GPCRs (50, 51). Overall, our results have unveiled a mechanism by which substitution of β-arrestin 2 by β-arrestin 1 in the absence of the normally beta cell predominant isoform is associated with acute defects on GLP-1R signalling, suggesting that the design of biased GLP-1R agonists away from recruitment of both β-arrestins should be favoured in order to elicit enhanced sustained GLP-1R responses without triggering acute deficits.

In the present study, we have also shown that reductions in acute GLP-1R cAMP are propagated towards acute deficits in calcium responses and insulin secretion in beta cell β-arrestin 2 KO islets, an effect that is eventually overturned during sustained exendin-4 stimulations, so that the net effect is an increased sustained *versus* acute insulin secretion responses in beta cell β-arrestin 2-deleted islets. Moreover, our *in vivo* beta cell connectivity analysis has unveiled that acute rises in connectivity triggered by GLP-1R agonism, previously observed in a separate study (52), only become sustained over time following β-arrestin 2 deletion, suggesting that GLP-1RAs biased away from β-arrestin recruitment might also trigger similar long-term improvements in intra-islet beta cell connectivity.

To further our knowledge of the molecular mechanisms underlying the prolonged signalling and *in vivo* glucoregulation observed in beta cell β-arrestin 2 absence, we have assessed islet GLP-1R trafficking patterns and, despite no differences in GLP-1R cell surface or internalisation levels in KO islets, we observed a significant reduction in active GLP-1Rs targeted to lysosomes, accompanied by concomitant reductions in receptor plasma membrane recycling and a redirection of stimulated receptors to the TGN in islets with beta cell-deleted β-arrestin 2. The lack of effect of β-arrestin 2 in GLP-1R internalisation, reminiscent of the behaviour of the GIPR in β-arrestin 2 knocked-down adipocytes (53), has previously been observed by our group using cell lines, and we now validate it here in primary islets. On the other hand, the reduced GLP-1R recycling result was unexpected, as it is opposite to that triggered by the biased compound exendin-phe1, which is associated with reduced recruitment of both β-arrestin isoforms to the GLP-1R (5). We nevertheless observed reduced GLP-1R lysosomal targeting, which does coincide with the reduced GLP-1R degradation triggered by exendin-phe1 (5), while at the same time detecting re-routing of the receptor to the TGN, a location from where it continued to signal. Overall, our results suggest that, in the absence of just β-arrestin 2, binding of the receptor to the remaining β-arrestin 1 results in a different trafficking phenotype than when binding to both β-arrestins is reduced with G protein-biased GLP-1RAs such as exendin-phe1.

Additionally, using an *in vivo* technique involving injection of near infrared-labelled exendin-4 derivative Ex-4-VT750 into mice, we could measure a significant loss of signal in cleared pancreata from control animals at 6-hours over 1-hour post-agonist injection that was no longer present in pancreata from β-arrestin 2 KO mice, suggesting potentially increased retention of pancreatic GLP-1R levels over prolonged agonist stimulation periods in the absence of β-arrestin 2, in agreement with the observed reduction in GLP-1R lysosomal targeting, and hence, presumably, GLP-1R degradation.

Most of our observations here are from *in vivo* or *ex vivo* islet experiments, but we have also created a rat insulinoma cell model with CRISPR/Cas9-deleted β-arrestin 2 to test our signalling and trafficking hypotheses in further detail. Using these cells, we have confirmed that β-arrestin 2 downregulation triggers reduced GLP-1R plasma membrane signalling efficacy without loss of endosomal signalling, accompanied by reduced acute (5-minute) fold increases in phospho-ERK1/2 and phospho-CREB in response to exendin-4 stimulation. Furthermore, as seen in islets, we have observed reduced targeting of active GLP-1Rs to recycling endosomes and lysosomes but increased interaction with the endosome-to-TGN marker Rab9, as well as increased TGN over global cAMP production, suggesting that, as for other GPCRs (54), TGN-rerouted GLP-1Rs in the absence of β-arrestin 2 are able to signal from this intracellular location. Overall, our observations point to a mechanism of signal prolongation in β-arrestin 2-deleted beta cells based in the avoidance of receptor degradation and access to TGN-based intracellular signalling platforms.

In parallel to studying GLP-1R behaviours, we have also performed a limited number of experiments on GIPR responses in beta cell β-arrestin 2 KO mice. Despite similar *in vivo* tendencies towards reduced sustained over acute glucose levels in KO animals treated with the stable GIP analogue D-Ala^2^-GIP, these did not quite reach statistical significance, suggesting a reduced effect of β-arrestin 2 on this receptor compared to that on the GLP-1R. Accordingly, acute insulin secretion in response to GIP showed only a tendency towards reduction in KO islets from chow-fed animals, although it did reach significance in islets from HFHS-fed mice, with the same tendency towards improvement in prolonged over acute exposures than for the GLP-1R. Also in agreement with a reduced influence of β-arrestin 2 on GIPR responses, no differences in plasma membrane or endosomal signalling were detected in GIP-stimulated β-arrestin 2 knockdown rat insulinoma cells.

Finally, and as β-arrestin 2 is known to be required for NEDD4 recruitment and ubiquitination of various GPCRs, including the β2 adrenergic (β2AR) (55), μ-opioid (MOR) and V2 vasopressin receptors (V2R) (56), in a mechanism that allows segregation of active receptors towards the degradative pathway following ubiquitin-specific binding to ESCRT machinery and GPCR lysosomal sorting (57), we have also investigated if a similar mechanism is in place for the GLP-1R which might explain the differences in lysosomal targeting and sustained signal prolongation in β-arrestin 2-deleted *versus* control beta cells, particularly as NEDD4 was one of the factors that we have recently identified as a direct GLP-1R interactor in a mass-spectrometry analysis of the GLP-1R beta cell interactome. Unexpectedly, we have found the GLP-1R to be constitutively ubiquitinated, with exendin-4 stimulation resulting in partial GLP-1R de-ubiquitination, a pattern also recently observed for the closely related glucagon receptor (58). In parallel, closely mimicking the ubiquitination results, NEDD4 was found constitutively recruited to the receptor, with this recruitment reduced following exendin-4 exposure. Importantly, there was a significant reduction in the overall level of GLP-1R ubiquitination and NEDD4 recruitment in cells without β-arrestin 2, which, surprisingly, was present even during vehicle conditions. Constitutive GPCR ubiquitination often functions to control receptor trafficking through the biosynthetic pathway (59), but we did not find any differences in cell surface GLP-1R levels in control *versus* β-arrestin 2 knocked down cells. Ubiquitination of certain GPCRs promotes their basal internalisation and lysosomal degradation, whereas deubiquitination often leads to recycling, switching the receptor’s fate and enhancing cellular resensitisation (59, 60). Our observations would fit with a complex pattern of GLP-1R ubiquitination/deubiquitination cycles that is disrupted in cells lacking β-arrestin 2, leading to divergent GLP-1R trafficking and signalling parameters.

In conclusion, we present here a comprehensive *in vivo*, *ex vivo* and *in vitro* assessment of the effects of knocking out β-arrestin 2 from primary beta cells on pharmacological GLP-1R responses, demonstrating reduced acute but enhanced long-term *in vivo* glucoregulation which correlates with acute *ex vivo* β-arrestin 1 and PDE4-dependent cAMP and downstream signalling defects but increased signal prolongation over time, a phenotype associated with a trafficking phenotype consisting on the diversion of active GLP-1Rs from lysosomal compartments towards the TGN and an overall reduction in GLP-1R ubiquitination and ubiquitin ligase NEDD4 recruitment to the receptor. Our study is the first to analyse in detail the consequences of selective β-arrestin 2 deletion from adult primary beta cells on GLP-1R behaviours. These data have important implications for the rational design of future GLP-1RAs with increased therapeutic windows.

Our study has several limitations, including difficulties in adjusting experimental conditions to translate *in vivo* acute and prolonged observations to the corresponding *ex vivo* trafficking and signalling assessments. Additionally, our trafficking studies are performed with fluorescently labelled exendin-4 and exendin-9 as a proxy for endogenous GLP-1R localisation, which is not entirely accurate. Better experiments could be performed in the future using animals genetically modified to express SNAP- or HALO-tagged GLP-1Rs from the endogenous locus. Finally, mechanistic studies in INS-1 832/3 cells will require validation in primary islets and experiments designed to establish causation from correlative data such as the effect of reduced GLP-1R ubiquitination in modulating GLP-1R post-endocytic trafficking and sustained signalling.

## Materials and Methods

### Animal studies

All *in vivo* procedures were approved by the UK Home Office under the Animals (Scientific Procedures) Act 1986 (Project Licences PA03F7F0F, P0A6474AE, and PP7151519 to Dr I. Leclerc, Dr B. Owen, and Dr A. Martinez-Sanchez, respectively) and from the local ethical committee (Animal Welfare and Ethics Review Board) at the Central Biological Services unit of Imperial College London. Animals were housed in groups of up to 5 adult mice in individually ventilated cages under controlled conditions (21-23°C; 12h-light:12h-dark cycles). *Ad libitum* access to standard chow diet was provided. For HFHS diet studies, animals were put on a 58 kcal % fat and sucrose diet (D12331, Research Diet, New Brunswick, NJ) *ad libitum* for the indicated time periods.

### Genomic DNA extraction and genotyping

Genotyping was carried out using ear samples collected from weaned animals. Extraction of genomic DNA was performed in an alkaline lysis buffer (25 mM NaOH, 0.2 mM EDTA pH 8.0 in dH_2_O) for 1 hour at 95°C, subsequently neutralised by the addition of 13 mM Tris-HCl, pH 7.4. The genomic DNA was used as a template for a PCR reaction with the appropriate primers (sequences provided in Table 1) and the Phire Green Hot Start II DNA polymerase (F124-L, Thermo Fisher Scientific, MA, USA). The PCR products were visualised on 1-2 % agarose gels using a Bio-Rad ChemiDoc imaging system.

**Table 1.**
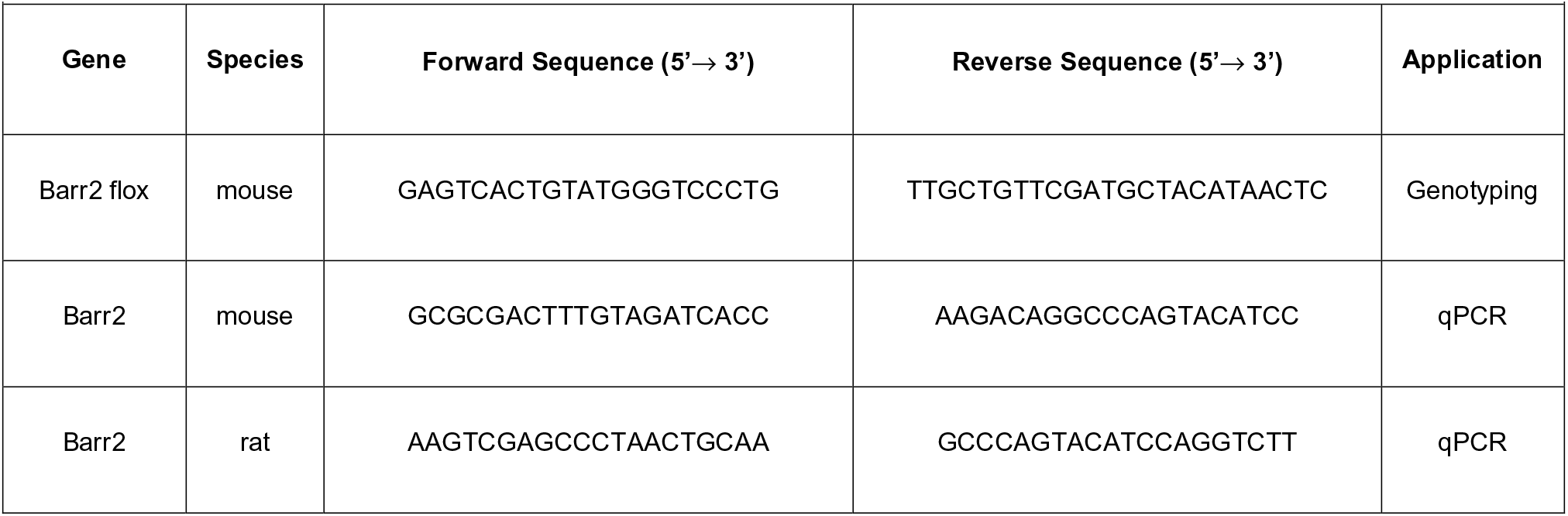
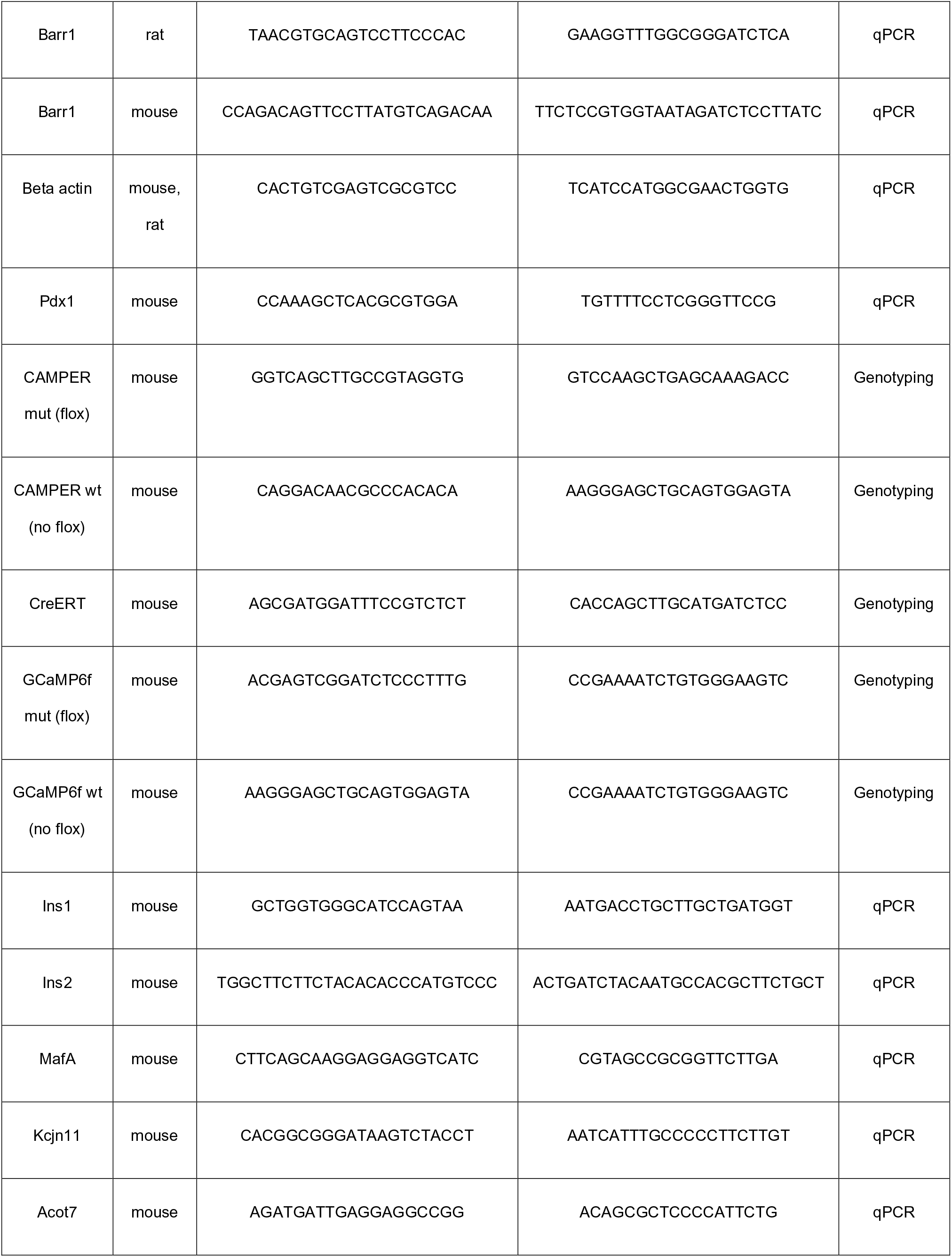
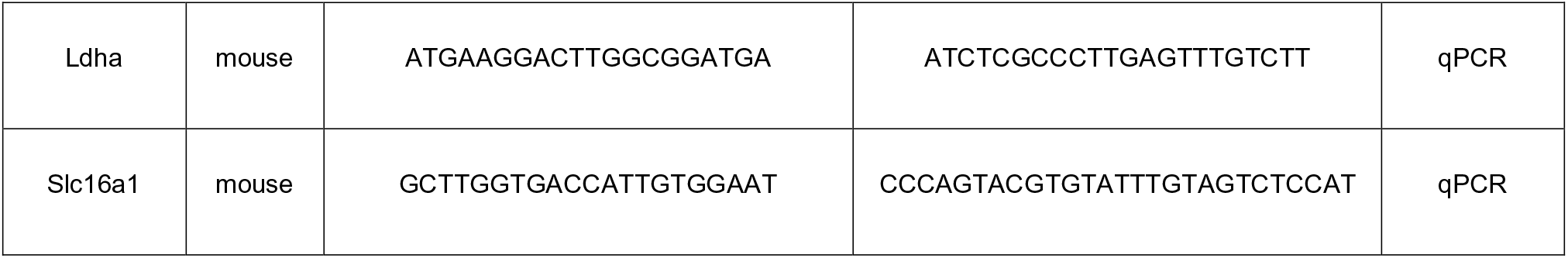
Primers used for genotyping PCR and qPCR.

### Generation of beta cell-selective β-arrestin 2 KO and control mice

Transgenic mouse models were generated on a C57BL/6 background using the Cre/Lox system. The Pdx1-Cre-ERT (Cre recombinase under the control of the *PDX1* promoter conjugated to a mutant oestrogen receptor sequence) mice were bred in-house, whereas the floxed β-arrestin 2 (Barr2) mice, in which exon 2 is flanked by loxP sites (Barr2 fl/fl), were kindly provided by Professor Marc Caron (Duke University, USA). Pdx1-Cre-ERT mice were crossed with Barr2 fl/fl mice in subsequent breeding pairs until mice hemizygous for the Pdx1-Cre-ERT and homozygous for Barr2 (Barr2 fl/fl) were obtained. These mice were then bred to produce Barr2 fl/fl mice with or without Pdx1-Cre-ERT. Barr2 fl/fl mice injected with tamoxifen were used as a control, since tamoxifen administration is a potential confounder (61, 62). Tamoxifen-treated Pdx1-Cre-ERT mice do not have altered beta cell function compared with wild-type littermates (63), thus, they were not used as an additional control. Tamoxifen (T5648, Sigma) dissolved at 20 mg/mL in corn oil (C8267, Sigma) was injected at a dose of 100 mg/kg intraperitoneally (i.p.) at 8 weeks of age for 5 consecutive days to induce Cre recombination and thus, beta cell-selective Barr2 KO mice and littermate controls were generated. Experiments were conducted at least 10 days after the last tamoxifen injection to allow for a sufficient tamoxifen washout period.

R26-Cre-ERT2 mice were kindly provided by Dr Tristan Rodriguez, Imperial College London, UK. This mouse line allows for ubiquitous tamoxifen-inducible Cre recombination, as a Cre-ERT2 cassette is inserted into the Rosa26 locus. These mice were crossed with Barr2 fl/fl mice until the generation of Barr2 fl/fl mice with or without expression of R26-Cre-ERT2. Tamoxifen administration as described above was used to produce inducible whole-body Barr2 KO and littermate control mice.

Mice that express the genetically encoded calcium indicator GCaMP6f Cre-dependently were used for *in vivo* calcium imaging experiments using islets implanted into the anterior chamber of the eye (37). GCaMP6f mice were bred in-house and crossed with Pdx1-Cre-ERT Barr2 fl/fl mice to generate GCaMP6f-Pdx1-Cre-ERT Barr2 fl/+ breeders. Subsequently, GCaMP6f-Pdx1-Cre-ERT Barr2 fl/fl and GCaMP6f-Pdx1-Cre-ERT Barr2 +/+ mice were obtained from the crossings and injected with tamoxifen to yield GCaMP6f+ beta cell Barr2 KO and control GCaMP6f+ Barr2 WT mice, respectively. In this case, the presence of the Cre recombinase is essential for GCaMP6f expression, as the conditional allele contains a loxP-flanked cassette.

Similarly, conditional expression of the biosensor ^T^EPAC^VV^ was utilised for imaging of cAMP dynamics: CAMPER reporter mice (29) were purchased from the Jackson Laboratory (Stock No: 032205) and crossed with Pdx1-Cre-ERT Barr2 fl/fl mice to generate CAMPER-Pdx1-Cre-ERT Barr2 fl/+ breeders. CAMPER-Pdx1-Cre-ERT Barr2 fl/fl and CAMPER-Pdx1-Cre-ERT Barr2 +/+ mice were obtained from the crossings and injected with tamoxifen resulting in CAMPER+ beta cell-specific Barr2 KO and control CAMPER+ Barr2 WT mice, respectively.

### Peptides

Peptides were produced by Wuxi AppTec Co. Ltd. (Shanghai, China), using standard solid phase peptide synthesis. Mass spectrometric confirmation of peptide identity and high-performance liquid chromatographic purity assessment were provided by the manufacturer (confirmed >90% purity). Tetramethylrhodamine (TMR)-labelled exendin-4 (exendin-4-TMR) and exendin-9 (exendin-9-TMR) have been described and validated before (64). VivoTag-750 conjugated exendin-4 (exendin-4-VT750), used for optical projection tomography (OPT) imaging, was synthesised and generously provided by Dr Sebastien Goudreau (ImmuPharma Group, Pessac, France).

### Intraperitoneal glucose tolerance tests (IPGTTs)

IPGTTs using 2 g/kg glucose were performed immediately and at the indicated time points after agonist injection (co-administered with glucose in the acute test) to examine acute and prolonged glycaemic responses to GLP-1R agonism. The animals were fasted for 2 hours before the tests starting at 8 am the morning of the experiment. Blood samples were analysed at 0, 10, 30, and 60 minutes after glucose ± agonist injection using a Contour glucometer (Bayer) and strips. The food was topped up at the end of the 6-hour IPGTT for agonists with shorter half-life (exendin-4, exendin-phe1, D-Ala^2^-GIP) or at the end of each IPGTT for agonists with longer half-life (tirzepatide, semaglutide).

### Measurement of in vivo plasma insulin levels

During selected IPGTTs, blood samples were collected at the indicated time points for plasma insulin analysis. Whole blood samples were collected in potassium EDTA cuvettes (Microvette CB 300, 16.444.100, Sarstedt) and centrifuged at 500 x g for 10 minutes at 4°C. The supernatant (plasma) was collected in fresh Eppendorf tubes and kept at 80°C until analysis with a mouse insulin ELISA kit (ultrasensitive mouse insulin ELISA kit, 90080, Crystal Chem), performed according to the manufacturer’s instructions with two technical replicates used per sample.

### Alpha and beta cell mass quantification

Pancreata from beta cell β-arrestin 2 KO and control littermates were dissected and fixed in 4% PFA for 24 hours. The tissues were washed twice in PBS and left in 70% ethanol until wax embedding. For each sample, three 5 μm-thick tissue sections separated by 300 μm were stained with guinea pig anti-insulin antibody (undiluted, Dako IR002; Alexa Fluor-488 secondary antibody, Invitrogen) and mouse anti-glucagon antibody (1:500, Sigma-Aldrich G2654; Alexa Fluor-568 secondary antibody, Invitrogen). Images were captured using a widefield Zeiss Axio Observer inverted microscope. Glucagon- and insulin-positive areas were determined as previously described (65) and expressed relative to total pancreas area imaged. The average islet size was approximated by adding the surface of beta cells and alpha cells for each islet per section, calculating the average, and producing the overall average for 3 separate sections per sample. Image J v1.53c was used for image analysis.

### Isolation and culture of pancreatic islets

For *ex vivo* islet experiments, pancreatic islets were isolated from appropriate KO and littermate control mice. Pancreata were infused via the common bile duct with RPMI-1640 medium (R8758, Sigma-Aldrich) containing 1 mg/mL collagenase from Clostridium histolyticum (S1745602, Nordmark Biochemicals), dissected, and incubated in a water bath at 37°C for 10 minutes. Islets were subsequently washed and purified using a Histopaque gradient (Histopaque-1119, 11191, Sigma-Aldrich, and Histopaque-1083, 10831, Sigma-Aldrich). Isolated islets were allowed to recover overnight at 37°C in 5% CO_2_ in RPMI-1640 supplemented with 10% v/v fetal bovine serum (FBS) (F7524, Sigma-Aldrich) and 1% v/v Penicillin/Streptomycin (P/S) solution (15070-063, Invitrogen).

### Implantation of islets into the anterior eye chamber

For chow diet calcium *in vivo* imaging, two 8-week-old female littermates with WT and KO genotypes for β-arrestin-2 (Barr2): one control (GCaMP6f heterozygous, Pdx1-Cre-ERT+ Barr2 +/+) and one KO (GCaMP6f heterozygous, Pdx1-Cre-ERT+ Barr2 fl/fl) were injected with tamoxifen for 5 consecutive days to induce GCaMP6f and Barr2 flox recombination. At 12 weeks of age, the animals were sacrificed, and pancreatic islets isolated. The day after the islet isolation, these were transplanted into the anterior chamber of the eye of six 12-week-old male WT C57BL/6J acceptors as in (37). Three of the acceptors received control and three KO donor islets. The success of the implantation was verified after 4 weeks, and imaging experiments carried out 8 weeks after the operation.

For calcium imaging under HFHS diet conditions two 8-week-old female littermates with WT and KO genotypes for β-arrestin-2 were concomitantly injected with tamoxifen and introduced to HFHS diet. Diet administration was continued for 8 weeks before animal sacrifice and islet isolation. The islets were implanted in the eyes of three 12-week-old female acceptors that had been on HFHS diet for 4 weeks. The acceptors received KO islets in their left eye and control islets in their right eye. The HFHS diet was continued throughout the study. Imaging experiments were carried out 4-5 weeks after the operation.

### In vivo calcium imaging

On the days of the imaging, the acceptor mice for chow diet experiments were injected i.p. with either vehicle (saline) or 10 nmol/kg exendin-4. For HFHS diet experiments, the mice were injected with 2 g/kg glucose with or without 10 nmol/kg exendin-4. All acceptor mice received both treatments using a cross-over study design with block randomization. Images were captured 30-minutes and 5- or 6-hours post-injection using a Nikon Eclipse Ti microscope with an ORCA-Flash 4.0 camera (Hamamatsu) and Metamorph software (Molecular Devices).

During the imaging experiments, general anaesthesia was induced using isoflurane, and time-lapse confocal microscopy performed using the 488 nm excitation channel for 181 frames with 800 msec exposure per frame. During the chow diet calcium imaging, mice injected with exendin-4 received intraperitoneal injections of 2 g/kg glucose (20% w/v glucose solution) to increase blood glucose levels shortly before image acquisition.

### Wave index assignment

Pancreatic beta cells are coupled such that healthy pulsatile insulin secretion is associated with pan-islet calcium oscillations or waves. We defined a wave index to objectively measure the proportion of an islet cross-section involved in calcium oscillatory activity. The wave index of each implanted islet was determined using Fiji: whole-islet mean intensity read-outs were obtained using a manual motion-correction macro developed by Mr. Stephen Rothery [National Heart and Lung Institute (NHLI) Facility for Imaging by Light Microscopy (FILM), Imperial College London]. Next, we determined the average whole islet calcium readout in an image sequence and multiplied this number by 1.2 to determine genuine calcium activity in islets. Using this value, the image sequences were thresholded to select cross-section areas with genuine calcium wave activity, expressed as a percentage relative to the whole islet cross-section area imaged. The highest percentage value in each image sequence was used to determine the islet’s wave index for that imaging session. Values were then corrected for the fact that, in implanted islets, approximately 20% of the imaged islet cross-section is covered by blood vessels. Using this method, we were able to determine the presence of four types of calcium wave activity, in line with earlier studies (38): islets where we observed activity over 0-25% of corrected area were characterized as type 1, if 26-50% of the islet cross-section was active these were typical of type 2, if 51-75% of the cross-section was active islets displayed type 3 activity and lastly, type 4 activity was assigned to islets where 76-100% of the cross-section was active.

### Waveform analysis

The wavelength, full width at half-maximum (FWHM), and amplitude was determined for the calcium traces of all islets imaged using MATLAB. Briefly, GCaMP6f fluorescence intensity traces were normalized to F_min_ and a cut-off value of 1.2 for genuine calcium activity was determined. Peaks in the data were determined using the derivative of the waveform for a given minimum peak height and distance between peaks. Frequency and wavelength were then calculated based upon peak location and value. Amplitude and FWHM were calculated as an average of all individual peak amplitude and FWHM for a given data set. Data without distinct peaking behaviour was discarded as noise.

### Connectivity analysis

For single cell connectivity analysis, individual beta cells were identified visually using the negative shadow of nuclei as a guidance for ROI placement. ROIs were subcellular and their XY coordinates and changes in mean intensity were measured. Using a MATLAB script, Pearson R correlation analysis of calcium time traces was performed. Calcium traces were normalised and smoothed using prospective time-points in the dataset. Pearson correlation between individual cell pairs was determined, excluding autocorrelation. An R-value of 0.25 was set as the threshold to signify a connection. Data were resampled using boot strapping to increase accuracy of findings (R-values with p<0.001 were deemed statistically significant). R-values were binned as follows: 0.25-0.50, 0.5-0.75 and 0.75-1, and considered to signify weak, medium, or strong connections, respectively. Cartesian line maps showing beta cell connectivity were generated based on cell XY coordinates and connections were assigned yellow, green, or red depending on strength of connections. Heat-map matrices show the R-values for each individual cell pair, with an R-value of 1 assigned for autocorrelation.

### Ex vivo calcium imaging

Imaging of whole-islet Ca^2+^ dynamics was performed 24 hours after isolation, essentially as previously described (52, 65). Islets from individual animals were pre-incubated for 1 hour in Krebs-Ringer Bicarbonate-HEPES (KRBH) buffer (140 mM NaCl, 3.6 mM KCl, 1.5 mM CaCl_2_, 0.5 mM MgSO_4_, 0.5 mM NaH_2_PO_4_, 2 mM NaHCO_3_, 10 mM HEPES, saturated with 95% O_2_/5% CO_2_; pH 7.4) containing 0.1% w/v BSA and 6 mM glucose (KRBH G6), and the Ca^2+^ responsive dye Cal-520, AM (AAT Bioquest). For chow diet experiments, islets were excited at 488 nm and images captured at 0.5 Hz using a Zeiss Axiovert microscope equipped with a 10X/0.5 numerical aperture objective and a Hamamatsu image-EM camera coupled to a Nipkow spinning-disk head (Yokogawa CSU-10). Volocity Software (PerkinElmer) provided a visualisation interface, while islets were maintained at 37°C on a heated stage constantly perifused with KRBH buffer containing G6 ± 100 nM exendin-4, 11 mM glucose, or 20 mM KCl. For HFHS diet experiments, islets were imaged with a Zeiss LSM-780 inverted confocal laser-scanning microscope in a 10X objective from the FILM Facility at Imperial College London. Treatments were manually added to the islet dishes by pipetting at the indicated time-points. To ensure that the islets remained stable, these were pre-encased into Matrigel (356231, Corning) and imaged on glass-bottom dishes (MatTek, P35G-1.5-10-C). Raw fluorescence intensity traces from whole islet ROIs were extracted using Image J v1.53c. Responses were plotted relative to the average fluorescence intensity per islet during the 6 mM glucose baseline period, before agonist addition.

### Islet GLP-1R trafficking experiments

*Ex vivo* islet GLP-1R trafficking experiments were carried out using intact islets treated in full media in 24-well suspension plates for the indicated time-points. The islets were loaded at the centre of a glass-bottom dish in RPMI without phenol red (32404014, Thermo Fisher Scientific). Z-stacks were obtained by confocal microscopy in a NHLI FILM Zeiss LSM-780 inverted confocal laser-scanning microscope and a 20X objective. The images were analysed using Image J v1.53c and processed using Z project with maximum intensity projections and the mean intensity of selected ROIs was measured.

### Ex vivo islet cAMP imaging

#### CAMPER islet FRET imaging

Intact islets from CAMPER-expressing mice (above) were used 24-48 hours following isolation. 10-20 islets from individual mice were encased into Matrigel on glass-bottom dishes and imaged for fluorescence resonance energy transfer (FRET) between CFP (donor) and YFP (acceptor) with CFP excitation and both CFP and YFP emission settings in a NHLI FILM Zeiss LSM-780 inverted confocal laser-scanning microscope and a 20X objective to capture time-lapse recordings with image acquisition every 6 seconds. The treatments were manually added by pipetting. Specifically, for acute cAMP studies, islets were imaged in KRBH buffer containing 0.1% w/v BSA and 6 mM glucose (KRBH G6) for 1 minute, then exendin-4 at 100 nM was added and imaged for 6 minutes before addition of isobutyl methylxanthine (IBMX) at 100 μM for the final 2 minutes of the acquisition. For acute rolipram experiments, islets were captured in KRBH G6 for 1 minute before addition of 100 nM exendin-4, 10 μM rolipram, or a combination of the two for 5 minutes. Finally, for overnight experiments, the islets were incubated in 24-well suspension plates in full media with 1 nM exendin-4 for 16 hours overnight, then washed for 30 minutes in KRBH G6, imaged for 2 minutes in this buffer, then 1 nM GLP-1 was added and finally, after 4 minutes, 100 μM IBMX was added and imaged for the final 2 minutes of the time-lapse. Raw intensity traces for YFP and CFP fluorescence were extracted from whole islet ROIs using Image J v1.53c and YFP/CFP ratios calculated for each ROI and time-point. Responses were plotted relative to the average fluorescence intensity per islet during the 6 mM glucose baseline period, before agonist addition.

#### Islet cAMP imaging with cADDis

Islets were infected with cADDis (Green Gi cADDis cAMP Assay Kit, Montana Molecular), a genetically encoded biosensor for cAMP packaged in a BacMam viral vector, following the manufacturer’s instructions, 24 hours after isolation. Infected islets were then imaged 24 hours post-infection. Islets were encased into Matrigel on glass-bottom dishes and imaged at 488 nm in a NHLI FILM Zeiss LSM-780 inverted confocal laser-scanning microscope and a 10X objective for time-lapse recordings with image acquisitions every 6 seconds. Treatments were manually added by pipetting. For acute cAMP studies, islets were imaged as above in KRBH buffer containing 0.1% w/v BSA and 6 mM glucose (KRBH G6) for 2 minutes to record the baseline, then exendin-4 at 100 nM was added and islets imaged for 5 minutes before addition of a mixture of 100 μM IBMX and 10 μM forskolin for the final 2 minutes of the acquisition. For overnight experiments, the islets were incubated in 24-well suspension plates in full media with 1 nM exendin-4 for 16 hours overnight, then washed for 30 minutes in KRBH G6, imaged for 2 minutes in this buffer, then 1 nM GLP-1 was added, and islets imaged for 5 minutes and finally, a mixture of 100 μM IBMX and 10 μM forskolin was added and imaged for the final 2 minutes of the acquisition. Raw intensity traces for GFP fluorescence were extracted from whole islet ROIs using Image J v1.53c and mean intensities calculated for each ROI and time-point. Responses were plotted relative to the average fluorescence intensity per islet during the 6 mM glucose baseline period, before agonist addition.

### β-arrestin 1 siRNA knockdown

For β-arrestin 1 (Barr1) siRNA studies, CAMPER primary mouse islets were dispersed by trituration in 0.05% trypsin/EDTA for 3 minutes at 37°C and seeded at the centre of poly-D-lysine-coated glass-bottom dishes. Dispersed islets were allowed to recover overnight before treatment with Accell mouse Arrb1 (109689) siRNA-SMARTpool (E-040976-00-0005, Horizon Discovery) or non-targeting control siRNA (D-001910-01-05, Horizon Discovery) according to the manufacturer’s instructions using the recommended siRNA buffer (B-002000-UB-100, Horizon Discovery) and serum-free delivery media (B-005000-100, Horizon Discovery). Studies took place 72 hours after siRNA treatment addition.

### Islet immunostaining for confocal co-localisation

For immunostaining and fluorescence confocal microscopy, exendin-4-TMR-treated islets were fixed using 4% PFA and stored in PBS. After a 10-minute permeabilisation with PBS containing 0.5 % v/v Triton X-100, the islets were washed once with PBS and incubated in blocking buffer (PBS, 0.1% v/v Tween-20, 1 % w/v goat serum, 1% BSA) for 30 minutes. Primary antibody against Lamp1 (1D4B, Developmental Studies Hybridoma Bank) or TGN38 (2F7.1, MA3-063, ThermoFisher Scientific) in blocking buffer was added overnight, while secondary anti-rat or anti-mouse Alexa Fluor 488 (A-11006, Thermo Fisher Scientific) was incubated for 30 minutes at room temperature. Islets were loaded onto glass-bottom dishes in PBS and z-stacks acquired by confocal microscopy with an Imperial College London NHLI FILM Zeiss LSM-780 inverted confocal laser-scanning microscope and a 63X/1.4 numerical aperture oil immersion objective for Lamp1 and an Imperial College London NHLI FILM Leica Stellaris 8 inverted confocal microscope and a 63X/1.4 numerical aperture oil immersion objective for TGN38. Images were analysed using Image J v1.53c. The Coloc 2 plugin was used for co-localisation analysis from maximum intensity projections.

### Optical projection tomography

Whole-body (R26-Cre-ERT2) β-arrestin 2 KO and control mice were injected i.p. with 100 nmol/kg of exendin-4-VivoTag 750 (exendin-4-VT750). After 1 hour or 6 hours, the mice were sacrificed using an overdose of anaesthetic (Euthatal solution for injection, Merial) and transcardially perfused with PBS followed by 4% PFA for perfusion-fixation. The pancreas and brain were dissected, washed once in PBS, and placed in 4% PFA for 1-6 hours. The tissues were subsequently optically cleared using the 3DISCO protocol (66). Dehydration was achieved by incubating in increasing concentrations (1x 50%, 1x 70%, 1x 80%, 3x 100%) of tetrahydrofuran (401757, Sigma) for 10-16 hours each time. Benzyl ether (108014, Sigma) was subsequently added for 16 hours.

The samples were imaged using an optical projection tomography microscope built by Dr James McGinty and Professor Paul French, Imperial College London. Scripts developed in MATLAB (MathWorks) by Dr McGinty were used for image acquisition, reconstruction, global scaling, and region segmentation. Quantification of object volumes and mean intensity was performed using 3D Objects Counter in Image J v1.53c. 3D images were visualised using Volocity software (Quorum Technologies Inc.).

### Cell culture

Cell lines were cultured in humidified incubators at 37°C in 5% CO_2_. INS-1 832/3 cells (67) (a gift from Dr Christopher Newgard, Duke University, USA) were grown in RPMI-1640 supplemented with 10% FBS, 1% P/S, 10 mM HEPES (H0887, Sigma), 1 mM sodium pyruvate (11360070, Thermo Fisher Scientific), and 0.05 mM 2-mercaptoethanol (M3148, Sigma). HEK293T cells were maintained in DMEM 4500 mg/L glucose (D6546, Sigma) with 10% FBS and 1% P/S. Cell lines were screened routinely for mycoplasma contamination.

### Generation of lentiviral CRISPR vectors and transduction of INS-1 832/3 cells

For knockdown (KD) of β-arrestin 2 in INS-1 832/3 cells, a lentiviral CRISPR approach was utilised. Two guide RNA (gRNA) sequences were cloned using a pScaffold-H1 (118152, Addgene) on a lentiCRISPR v2 backbone (52961, Addgene), using a protocol adapted from (68). The gRNA sequences were: 5’-GAAGTCGAGCCCTAACTGCA-3’ and 5’- ACCGGTATTTGAAGCCTCTT-3’ (reverse complement). The backbone vector was pre-digested with FastDigest Esp3I (FD0454, Thermo Fisher Scientific) for 2 hours at 37°C and purified using Monarch® DNA Gel Extraction Kit (T1020S, New England Biolabs) before digestion-ligation with the gRNA-pScaffold-H1 using FastDigest Esp3I and T7 Ligase (M0318L, New England Biolabs).

To produce lentiviral particles, HEK293T cells were co-transfected with the gRNA lentiviral vector (or the vector without gRNAs - control empty vector), plus the packaging (psPAX2) and envelope (pMD2.G) vectors using a calcium phosphate protocol. Viral supernatants were harvested 48- and 72-hours post-transfection, filtered using a 0.45 μm Millex-HV filter, and concentrated by 20% sucrose gradient ultracentrifugation in an Optima XPN-100 ultracentrifuge at 26,000 rpm at 4°C for 2 hours in a SW32 Ti swinging bucket rotor (Beckman-Coulter). Viral particles were resuspended in PBS and stored at −80°C.

INS-1 832/3 cells were transduced with appropriate amounts of lentiviruses followed by addition of puromycin (4 μg/ml) 72 hours post-transfection. Transduction was performed with the control empty vector (INS-1 832/3 EV) or the vector with gRNAs 1 and 2 (INS-1 832/3 g1-2) to generate β-arrestin 2 KDs. The puromycin was replaced every 2-3 days for a total of 2 weeks to induce the selection of transduced cells. The surviving selected cells were subsequently cultured in full media in the absence of puromycin and tested for β-arrestin 2 gene expression using qPCR.

### Ex vivo insulin secretion assays

Isolated islets used for insulin secretion assays were treated in 24-well non adherent plates. Ten islets were used per well and three technical replicates used per condition. For acute studies, islets were pre-incubated for 1 hour in KRBH buffer (described in *in vitro* calcium imaging) containing 1% w/v BSA (10775835001, Roche) and 3 mM glucose before incubation with 11 mM glucose ± agonists in KRBH in a shaking 37°C water-bath (80 rpm) for 1 hour. For overnight studies, pre-incubation was carried out in RPMI-1640 media containing FBS, P/S and 3 mM glucose, followed by treatment with media containing 11 mM glucose ± agonists for 16 hours. At the treatment end, supernatants containing the secreted insulin were collected, centrifuged at 1,000 x g for 5 minutes, and transferred to fresh tubes. To determine total insulin content, islets were lysed using acidic ethanol (75% v/v ethanol, 1.5 mM HCl). The lysates were sonicated 3 x 10 seconds in a water bath and centrifuged at 10,000 x g for 10 minutes, and the supernatants collected. The samples were stored at −20°C until the insulin concentration was determined using an Insulin Ultra-Sensitive Homogeneous Time Resolved Fluorescence (HTRF) Assay kit (62IN2PEG, Cisbio, Codolet, France) according to the manufacturer’s instructions. GraphPad Prism 9 was used for the generation of the standard curve and sample concentration extrapolation. The total insulin content was calculated by adding the secreted insulin to the insulin content of the lysates.

### HTRF cAMP assays

Intact primary mouse islets were dispersed by trituration in 0.05% trypsin/EDTA for 3 minutes at 37°C. After stimulation, dispersed islets were lysed and cAMP assayed by HTRF immunoassay (cAMP Dynamic 2, 62AM4PEB, Cisbio, Codolet, France).

### Transfection of plasmid DNA

Transient transfection of cell lines with plasmid DNA was achieved with Lipofectamine 2000 Transfection Reagent (11668027, Thermo Fisher Scientific) according to the manufacturer’s instructions. Briefly, appropriate amounts of DNA were incubated for 5 minutes in Opti-MEM reduced serum medium (31985070, Thermo Fisher Scientific) and gently mixed with an equal volume of Lipofectamine 2000 in Opti-MEM. After 20 minutes, the mix was added dropwise to cells in culture medium without P/S and incubated for 4-6 hours. The medium was then removed, and full culture medium added. Assays were performed 24 hours or 48 hours post-transfection depending on the experiment.

### RNA extraction and qPCR

Samples were lysed in TRIzol Reagent (15596-018, Invitrogen) and briefly vortexed for homogenisation. Phase separation was achieved by chloroform (C2432, Sigma) addition and the upper aqueous phase collected. RNA was recovered by overnight precipitation with isopropanol (11388461, Thermo Fisher Scientific). Following cDNA synthesis using MultiScribe™ Reverse Transcriptase (4311235, Thermo Fisher Scientific) according to the manufacturer’s instructions, qPCR was performed using SYBR Green Technology (Fast SYBR Green Master Mix, 4385616, Invitrogen) on an Applied Biosystems™ 7500 Real-Time PCR system. The data was analysed using the 2^(-ΔΔCt)^ method (69). A list of primer sequences is provided in Table 1.

### Cyclic AMP FRET imaging using global and TGN-targeted ^T^EPAC^VV^

The global ^T^EPAC^VV^ construct was kindly provided by Professor Kees Jalink, the Netherlands Cancer Institute, Netherlands (70). The Trans-Golgi network (TGN)-targeted version was made *in house* by in-frame cloning of the GRIP domain of GolginA1 (pCMV6-KL5-GolginA1, Origene) at the C-terminal end of ^T^EPAC^VV^. INS-1 832/3 EV and g1-2 cells were plated in 48-wells and transfected with 0.5 µg plasmid DNA. After 24 hours, cells were trypsinised and seeded on glass-bottom dishes. 48 hours post-transfection, cells were imaged in RPMI without phenol red (32404014, Thermo Fisher Scientific) with a NHLI FILM Zeiss LSM-780 inverted confocal laser-scanning microscope using a 20X objective to capture a time-lapse recording of CFP/YFP FRET as described for CAMPER islets with image acquisition every 6 seconds and treatments manually added by pipetting. Cells were imaged for 1 minute to record a baseline, then exendin-4 at 100 nM was added and imaged for 4 minutes before addition of 100 μM IBMX and 10 μM forskolin for the final 2 minutes of acquisition. Raw intensity traces for YFP and CFP fluorescence were extracted from individual cell ROIs using Image J v1.53c and YFP/CFP ratios calculated for each ROI and time-point. Responses were plotted relative to the average fluorescence intensity per islet during the 6 mM glucose baseline period, before agonist addition.

### NanoBiT complementation and NanoBRET assays

For Nb37-based bystander complementation assays, the plasmids used were a gift from Prof Asuka Inoue, Tohoku University, Japan (40). Nb37 cDNA (synthesised by GenScript with codon optimisation) was C-terminally fused to SmBiT with a 15-amino-acid flexible linker (GGSGGGGSGGSSSGGG), and the resulting construct referred to as Nb37-SmBiT. The C-terminal KRAS CAAX motif (SSSGGGKKKKKKSKTKCVIM) was N-terminally fused with LgBiT (LgBiT-CAAX). The Endofin FYVE domain (amino-acid region Gln739-Lys806) was C-terminally fused with LgBiT (Endofin-LgBiT). Gαs (human, short isoform), Gβ1 (human), Gγ2 (human), and RIC8B (human, isoform2) plasmids were inserted into pcDNA 3.1 or pCAGGS expression vectors. INS-1 832/3 EV or g1-2 cells were seeded in 6-well plates and co-transfected with 0.2 μg SNAP-GLP-1R or SNAP-GIPR, 0.5 μg Gαs, Gβ1, and Gγ2, 0.1 μg RIC8B, 0.1 μg CAAX-LgBiT or 0.5 μg Endofin-LgBiT with 0.1 μg or 0.5 μg Nb37-SmBiT (1:1 ratio), respectively.

For NanoBRET localisation assays, cells were seeded in 12-well plates and co-transfected with SNAP-GLP-1R-NLuc (generated in-house) and KRAS-, Rab4-, Rab5-, Rab9-, or Rab11-Venus (gifts from Prof Nevin Lambert, Augusta University, USA, and Prof Kevin Pfleger, The University of Western Australia, Australia). For KRAS-, Rab5-, and Rab11-Venus, 0.5 µg were co-transfected with 0.5 µg SNAP-GLP-1R-NLuc, while for Rab4- and Rab9-Venus, 0.25 µg were co-transfected with 0.1 µg SNAP-GLP-1R-NLuc. 24 hours after transfection, cells were detached, resuspended in Nano-Glo® Live Cell Reagent (N2011, Promega) with furimazine (1:20 dilution) and seeded into white 96-well half area plates. Luminescence was recorded at 37°C in a Flexstation 3 plate reader, with total luminescent signal used for NanoBiT assays and dual wavelength acquisition (460 and 535 nm) for NanoBRET assays. A 5-minute baseline recording was followed by agonist addition and serial measurements over 30 minutes. Readings were taken every 30 seconds and normalised to well baseline and then, average vehicle-induced signal was subtracted to establish the agonist-induced effect. Areas under the curve (AUC) were calculated for each agonist concentration and fitted to four-parameter curves using GraphPad Prism 9.0.

### Binding kinetic assay in primary islets

Islets were pre-incubated at 37°C with 1 µM exendin-4 in RPMI-1640 (with FBS and P/S) for 2 hours prior to imaging to saturate receptor binding to generate a negative control for TMR uptake, with exendin-4 maintained for the duration of imaging. These, along with islets not pre-incubated with exendin-4, were incubated with a metabolic inhibitor cocktail (20 mM 2-deoxyglucose, 10 mM NaN_3_) to inhibit ATP-dependent endocytosis, as previously described (71), for 30 minutes prior to imaging. The metabolic inhibitors were maintained for the duration of the imaging.

Approximately 20 islets were then encased onto Matrigel in glass-bottom dishes and imaged in imaging medium (RPMI-1640 without phenol red, containing metabolic inhibitors) with a Zeiss LSM-780 inverted confocal laser-scanning microscope with a 10X objective from the FILM Facility at Imperial College London. TMR fluorescence was recorded every 2 seconds for 1 minute baseline, followed by 100 nM exendin-4-TMR addition in imaging medium and recording for a further 15 minutes. Curve fitting to an exponential plateau was performed to calculate binding kinetic parameters in Prism 9.0.

### Generation of stable SNAP-GLP-1R-expressing beta cell sublines

Five million INS-1 832/3 EV or g1-2 cells were seeded into 10-cm adherent dishes. Each dish was transfected with 9 µg of the SNAP-GLP-1R plasmid (Cisbio) using Lipofectamine 2000 according to the manufacturer’s protocol. Forty-eight hours later, 1 mg/mL G418 was added to each dish to select for SNAP-GLP-1R-positive cells. The surviving cells were allowed to proliferate. Once the 10-cm dishes reached >80% confluence, cells were labelled in suspension with SNAP-Surface 549 (New England Biolabs) for 30 minutes at 37°C and FACS-sorted to select for SNAP-GLP-1R-expressing ones. Sorted cells were then cultured and maintained in 1 mg/mL G418 to preserve SNAP-GLP-1R expression.

### Immunoprecipitation assays

For NEDD4 co-immunoprecipitation, INS-1 832/3 EV or g1-2 SNAP-GLP-1R cells were transfected with the pCI HA NEDD4 construct (a gift from Prof. Joan Massague, Addgene plasmid #27002) 24 hours before the experiment. Two million INS-1 832/3 EV or g1-2 SNAP-GLP-1R cells were seeded onto cell culture treated 6-well plates and allowed to attach overnight. The next day, the cells were treated with vehicle or 1 µM exendin-4 in RPMI-1640 (containing FBS, P/S and additional HEPES, sodium pyruvate and 2-mercaptoethanol) for 10 minutes at 37°C. Following stimulation, cells were washed 1X in ice-cold PBS and lysed in ice-cold 1X TBS (50 mM Tris-HCl pH 7.4, 150 mM NaCl) supplemented with 1 mM EDTA, 1% Triton X-100, and protease and phosphatase inhibitor cocktails and lysates placed on a rocker for 15 minutes at 4°C. Next, lysates were incubated overnight under rotation at 4°C with ANTI-FLAG® M2 Affinity Gel beads (Merck) to pull down FLAG-containing SNAP-GLP-1R, and pull-downs were performed according to the manufacturer’s protocol.

Following the pull-down, the beads were resuspended in 1X TBS and 2X urea loading buffer (20 mM Tris-HCl pH 6.8, 5% SDS, 8 M urea, 100 mM dithiothreitol, 0.02% bromophenol blue) 1:1 and incubated at 37°C for 10 minutes to separate pulled down proteins from beads. Samples were spun at 5000 x g for 30 seconds and the supernatant resolved in acrylamide gels, electroblotted onto polyvinylidene fluoride (PVDF) membranes, incubated with the indicated antibodies and developed as detailed in the Western blotting section. Specific band densities were quantified with Image J v1.53c. The details for the primary antibodies for ubiquitin and HA-tag and respective secondary antibodies are provided in Table 2.

**Table 2.** Primary and secondary antibodies used for Western blotting.

| 1° Ab | Manufacturer | Cat. No. | Dil. | Time, T° | 2° Ab | Manufacturer | Cat. No. | Dil. | Time, T° |
| --- | --- | --- | --- | --- | --- | --- | --- | --- | --- |
| ERK 1/2 | Cell Signaling Technology | 9102S | 1:1000 | 16 h, 4°C | Rabbit | Abcam | ab205718 | 1:5000 | 1 h, RT |
| pERK1/2 | Cell Signaling Technology | 9101S | 1:1000 | 16 h, 4°C | Rabbit | Abcam | ab205718 | 1:5000 | 1 h, RT |
| CREB | Cell Signaling Technology | 9197S | 1:1000 | 16 h, 4°C | Rabbit | Abcam | ab205718 | 1:5000 | 1 h, RT |
| pCREB | Cell Signaling Technology | 9198S | 1:1000 | 16 h, 4°C | Rabbit | Abcam | ab205718 | 1:5000 | 1 h, RT |
| SNAP-tag | New England Biolabs | P9310S | 1:500 | 16 h, 4°C | Rabbit | Abcam | ab205718 | 1:5000 | 1 h, RT |
| Ubiquitin | Santa Cruz Biotechnology | sc-8017 | 1:1000 | 16 h, 4°C | Mouse | Abcam | Ab6728 | 1:5000 | 1 h, RT |
| HA-tag | Sigma | H3663 | 1:1000 | 16 h, 4°C | Mouse | Abcam | Ab6728 | 1:5000 | 1 h, RT |

### Protein extraction and Western blotting

Protein extraction was performed by lysing samples with Tris/NaCl/EDTA buffer (100 mM NaCl, 50 mM Tris-HCl, 1 mM EDTA, 0.1% BSA, pH 8.6). The samples were sonicated in a water bath sonicator 3X for 10 seconds. 2X urea loading buffer was added 1:1 and samples were incubated at 37°C for 10 minutes before resolving using SDS-PAGE (10% acrylamide gels). Protein transfer to PVDF membranes (Immobilon-P, 0.45 µm pore size, IPVH00010, Merck) was achieved using a wet transfer system (Bio-Rad). The membranes were incubated with appropriate primary and secondary antibodies listed in Table 2 in 5% milk and developed using the Clarity Western enhanced chemiluminescence substrate system (1705060, Bio-Rad) in a Xograph Compact X5 processor. Specific band densities were quantified using Image J v1.53c.

### Cell labelling for confocal co-localisation

INS-1 832/3 EV and g1-2 cells transiently expressing the SNAP-GLP-1R were labelled at 37°C with 1 μM of SNAP-Surface 649 fluorescent substrate (S9159S, New England Biolabs) in full media prior to treatment with 100 nM exendin-4 or vehicle for 3 hours. 5 minutes before the end of the latter incubation period, 1 μM LysoTracker Red DND-99 (L7528, Thermo Fisher Scientific) was added. The cells were washed in PBS and fixed in 4% paraformaldehyde, mounted in Prolong Diamond antifade reagent with 4,6-diamidino-2-pM phenylindole (Life Technologies), and imaged by confocal microscopy with an Imperial College London NHLI FILM Zeiss LSM-780 inverted confocal laser-scanning microscope and a 63X/1.4 numerical aperture oil immersion objective equipped with Zen software (ZEN 2.3 SP1, black, 64-bit, Carl Zeiss). Images were analysed using Image J v1.53c. The Coloc 2 plugin was used for co-localisation analysis.

### Time-lapse β-arrestin 2 – GLP-1R co-localisation by spinning disk microscopy

INS-1 832/3 cells stably expressing SNAP-GLP-1R were transiently transfected with a β-arrestin 2-GFP construct. 24 hours after transfection, cells were seeded onto glass bottom MatTek dishes and left to adhere overnight. Cells were labelled in full media with SNAP-Surface 549 (New England Biolabs) for 30 minutes at 37°C and imaged in RPMI without phenol red in a Nikon Eclipse Ti spinning disk confocal microscope with an ORCA-Flash 4.0 camera (Hamamatsu) and Metamorph software (Molecular Devices). Time-lapse images of green and red fluorescence were acquired every 15 seconds for an initial 5-minute baseline prior to addition of 100 nM exendin-4 and further imaging for 10 minutes.

### Electron microscopy

Islets extracted from chow diet-fed beta cell β-arrestin 2 KO and control littermates were fixed in EM-grade 2% PFA + 2% glutaraldehyde mix overnight in 0.1 M cacodylate buffer and conventional EM was performed as described (5). Briefly, following fixation, islets were post-fixed with osmium tetroxide, encased in 1% agarose, processed for EM, and embedded in Epon resin which was then polymerized at 60°C overnight. 70□nm-thick sections were cut with a diamond knife (DiATOME) in a Leica Ultracut UCT ultramicrotome before examination on an FEI Tecnai T12 Twin TEM. Images were acquired in a charge-coupled device camera (F216, TVIPS), and processed in Image J v1.53c.

### Statistical analyses

For single cell connectivity analysis, statistical differences were evaluated using paired or unpaired Student’s t-test in MATLAB (MathWorks). All other data analyses and graph generation were performed with GraphPad Prism 9.0. The statistical tests used are indicated in the corresponding figure legends. The number of replicates for comparisons represents biological replicates. Technical replicates within biological replicates were averaged prior to statistical tests.

Data are represented as mean ± SEM, unless otherwise stated. The p-value threshold for statistical significance was set at 0.05.

## Supporting information

Graphical Abstract

Barr2-GFP SNAP-GLP-1R

## Supplemental Figure Legends

**Supplemental Figure 1. GLP-1R agonist responses in lean adult beta cell-selective** β-arrestin 2 KO *vs* control mice – extra data. IPGTTs (2 g/kg glucose i.p.) were performed concurrently with, or 6 h after, i.p. administration of agonists or vehicle (saline). (**A**) Diagram depicting the tamoxifen-induced adult beta cell-selective β-arrestin 2 KO mechanism using the Cre-lox system. (**B**, **C**) Weight (B), and fasting and fed glycaemia (C) of lean adult beta cell-selective β-arrestin 2 (β-Barr2) KO *vs* control male and female mice (n = 6-9 / genotype and sex, age: 20-24 weeks). (**D**) Representative EM images depicting islet ultrastructure and quantification of insulin granule densities in islets isolated from lean adult β-Barr2 KO *vs* control mice on chow diet (n = 5). (**E**) Glucose curves for vehicle or exendin-4 (Ex-4) administration at 0.1 and 10 nmol/kg in lean, male mice (n = 8 / genotype, age: 12-16 weeks). (**F**) Glucose curves for vehicle or Ex-4 administration at 0.1 and 10 nmol/kg in lean, female mice (n = 9 / genotype, age: 12-16 weeks). (**G**) Absolute and fold-change *vs* vehicle values for 10-min plasma insulin concentrations during IPGTTs (2 g/kg glucose i.p.) performed concurrently with, or 6 h after administration of vehicle or 1 nmol/kg Ex-4 in lean female mice (n = 7-9 / genotype, age: 12-16 weeks). Comparisons were performed using unpaired t-test or two-way ANOVA with Sidak’s *post hoc* tests. Data are presented as mean ± SEM.

**Supplemental Figure 2. GIPR agonist responses in adult beta cell-selective** β-arrestin 2 KO *vs* control mice and GLP-1R agonist responses in whole body β-arrestin 2 KO *vs* control mice. IPGTTs (2 g/kg glucose i.p.) performed concurrently with or 6 h after i.p. administration of agonists or vehicle (saline). (**A**, **B**) Glucose curves (A) and corresponding ΔAUCs (6h-0h) (B) for vehicle or D-Ala^2^-GIP administration at 10 and 40 nmol/kg in lean male mice (n = 8 / genotype, age: 18-26 weeks). (**C**, **D**) Glucose curves (C) and corresponding ΔAUCs (6h-0h) (D) for vehicle or D-Ala^2^-GIP administration at 10 and 40 nmol/kg in lean female mice (n = 8-9 / genotype, age: 18-26 weeks). (**E**, **F**) Glucose curves (E) and corresponding ΔAUCs (6h-0h) (F) for vehicle or Ex-4 administration at 0.1, 1, and 10 nmol/kg in tamoxifen-inducible whole-body (R26-Cre-ERT) Barr2 KO and control lean males (n = 8 / genotype, age: 12-16 weeks). (**G**, **H**) Glucose curves (G) and corresponding ΔAUCs (6h-0h) (H) for vehicle or Ex-4 administration at 0.1, 1, and 10 nmol/kg in tamoxifen-inducible whole-body (R26-Cre-ERT) Barr2 KO and control lean females (n = 8 / genotype, age: 12-16 weeks). Comparisons were performed using two-way ANOVA or mixed-effects model with Sidak’s *post hoc* tests. Data are presented as mean ± SEM.

**Supplemental Figure 3. HFHS-fed adult beta cell-selective** β**-arrestin 2 KO *vs* control mice – extra data.** (**A**) Weekly weights of male and female mice after HFHS diet initiation (n = 7-9 / genotype and sex). (**B**) Fasting and fed glycaemia of HSHF diet-fed male and female mice (n = 5-7 / genotype and sex, duration of HSHF diet: 10-16 weeks). (**C**) Representative images of pancreatic sections from adult beta cell-selective β-arrestin 2 (β-Barr2) KO *vs* control mice on HFHS diet depicting islets with nuclei stained with DAPI (blue) and co-stained for insulin (green) and glucagon (red). (**D**) Quantifications of beta and alpha cell mass, alpha/beta cell mass ratio and average islet sizes in adult β-Barr2 KO *vs* control mice on HFHS diet (n = 5-6 / group). (**E**) Relative gene expression of selected beta cell-enriched and -disallowed genes in control *vs* adult β-Barr2 KO mice on chow (n = 6) or HFHS diet (n = 4). (**F**) Absolute and fold-change *vs* vehicle values for 10-min plasma insulin concentrations during IPGTTs (2 g/kg glucose i.p.) performed concurrently with or 6 h after administration of vehicle or 1 nmol/kg Ex-4 in mixed male and female mice on HFHS diet (n = 7 / genotype, duration of HFHS diet: 10-16 weeks). Comparisons were made using unpaired t-tests or one or two-way ANOVA with Sidak’s *post hoc* tests. *p<0.05, **p<0.01, ***p<0.001, ****p<0.0001 *vs* control group. Data are presented as mean ± SEM.

**Supplemental Figure 4. Extra *ex vivo* signalling data from adult beta cell-selective** β**-arrestin 2 KO *vs* control islets.** (**A**, **B**) cAMP dose-response curves and corresponding Emax and logEC50 values in dispersed islets isolated from adult beta cell-selective β-arrestin 2 (β-Barr2) KO and control mice on (A) chow diet (n= 5-6 / group) or (B) HFHS diet (n= 4-5 / group). (**C**) cAMP responses over time from cADDis-infected beta cell-specific Barr2 KO *vs* control islets in response to 100 nM Ex-4 followed by 100 μM IBMX + 10 μM forskolin (FSK); AUCs calculated for the Ex-4 treatment period (n = 4 / genotype). (**D**) cAMP responses over time from cADDis-infected β-Barr2 KO *vs* control islets pre-treated with 1 nM Ex-4 for 16 h (overnight) in response to 1 nM GLP-1 stimulation followed by 100 μM IBMX + 10 μM FSK; AUCs calculated for the GLP-1 treatment period (n = 4 / genotype). (**E**) Percentage of islets within each wave activity index category for chow and HFHS diet animals (n = 6-13 islets / condition). (**F**, **G**) Calcium wave characteristics: amplitude, wavelength, and full width at half maximum (FWHM) for beta cell-selective β-arrestin 2 (β-Barr2) KO *vs* control GCaMP6f islets from donors implanted in the anterior chamber of the eye from (F) chow and (G) HFSH diet experiments (n = 0-13 islets / condition). (**H**) Percentage of connectivity for β-Barr2 KO *vs* control GCaMP6f islets from chow diet animals implanted in the anterior chamber of the eye of chow diet-fed WT animals, which received Ex-4 10 nmol/kg or saline (vehicle) i.p. (n = 2-11 islets / condition). (**I**, **J**) Insulin secretion assays from control *vs* β-Barr2 KO islets treated with 100 nM Ex-4 or 100 nM GIP (fold over 11 mM glucose vehicle levels) for 1 h (acute) or 16 h (overnight) isolated from mice on (I) chow diet (n = 8 / genotype) or (J) HFHS diet (n = 6 / genotype). Comparisons were made with t-tests or two-way ANOVA with Sidak’s *post hoc* tests. *p<0.05, **p<0.01 *vs* control group. Data are presented as mean ± SEM.

**Supplemental Figure 5. Extra *ex vivo* trafficking data from adult beta cell-selective** β**-arrestin 2 KO *vs* control islets.** (**A**) Ex-4-TMR fluorescence at 1 h in β-Barr2 KO *vs* control islets treated or not with acetic acid wash for 5 min before imaging for animals on chow (n = 6 / genotype) or HFHS diet (n = 4 / genotype). (**B**) Quantification of TMR fluorescence in β-Barr2 KO *vs* control islets pre-treated with vehicle or 100 nM Ex-4 for 1 h and then washed and treated with 100 nM Ex-4-TMR for 3 h for animals on chow (n = 6 / genotype) or HFHS diet (n = 4 / genotype). (**C**) TMR fluorescence in β-Barr2 KO *vs* control islets pre-treated with vehicle or 100 nM Ex-4 for 1 h and then washed and treated with 100 nM Ex-4-TMR for 6 h in chow diet animals (n = 9 / genotype). Comparisons were made with two-way ANOVA with Sidak’s *post hoc* tests. Data are presented as mean ± SEM.

**Supplemental Figure 6. β-arrestin 2 recruitment dynamics in INS-1 832/3 SNAP-GLP-1R cells.** (**A**) Time-lapse spinning disk imaging of SNAP-GLP-1R (red) and β-arrestin 2-GFP (green) dynamic profiles in INS-1 832/3 SNAP-GLP-1R cells in response to 100 nM Ex-4. Selected time frames are depicted for the green, red, and merged channels. **(B**) High magnification image for the 5-min stimulation time-point from (A). Areas of SNAP-GLP-1R - β- arrestin 2-GFP co-localisation are indicated with white arrows.

**Supplemental Figure 7. Extra data from β-arrestin 2 KD *vs* control INS-1 832/3 cell lines.** (**A**) D-Ala^2^GIP dose-response AUC curves for Nb37-SmBiT and CAAX-LgBiT (plasma membrane) or Endofin-LgBiT (endosomal) signal complementation assays (n = 5). (**B**) LogEC50 and Emax values calculated from (A). (**C**) Quantification of pERK1/2 over total ERK1/2, and pCREB over total CREB using densitometry analysis (n = 4 for pERK1/2 and n = 3 for pCREB). (**C**) Endogenous surface GLP-1R receptor levels in untreated β-arrestin 2 (Barr2) KD *vs* control INS-1 832/3 cells quantified by labelling for 1 h with 1 µM Ex-9-TMR (n = 6). (**E**) Schematic representation of the principle of NanoBRET-based subcellular localisation assays. (**F**) Quantification of SNAP-GLP-1R deubiquitination in response to 10 min of 100 nM Ex-4 exposure in INS-1 832/3 SNAP-GLP-1R control and Barr2 KD cells. Data from Figure 7F normalised to vehicle conditions for each cell type. (**G**) Quantification of loss of HA-NEDD4 - SNAP-GLP-1R interaction in response to 10 min of 100 nM Ex-4 exposure in INS-1 832/3 SNAP-GLP-1R Barr2 KD and control cells. Data from Figure 7G normalised to vehicle conditions for each cell type. Comparisons were made with ratio-t-tests and two-way ANOVA with Sidak’s *post hoc* tests. *p<0.05, **p<0.01, ***p<0.001 *vs* control group. Data are presented as mean ± SEM.

## Acknowledgements

This work was supported by MRC grant number MR/R010676/1 to A.T., B.J. and G.A.R., and by UKRI COVID-19 Grant Extension Allocation (coA) to A.T. A.T. also acknowledges further support from the MRC, EFSD, Diabetes UK, Eli Lilly, the Commonwealth, and the Integrated Biological Imaging Network (IBIN). S.B. was supported by an Early Career Grant from the Society for Endocrinology. GAR was additionally supported by Wellcome Trust Senior Investigator Awards (098424AIA, 212625/Z/18/Z), MRC Programme grants (MR/R022259/1, MR/J0003042/1, MR/L020149/1), an Experimental Challenge Grant (DIVA, MR/L02036X/1), an MRC grant (MR/N00275X/1), and Diabetes UK (BDA16/0005485). The authors thank Mr Stephen Rothery from the National Heart and Lung Institute (NHLI) Facility for Imaging by Light Microscopy (FILM), Imperial College London, for technical assistance and help with data analysis, Dr Pauline Chabosseau for providing a macro for beta and alpha cell mass quantification in Image J and Dr Sebastien Goudreau from ImmuPharma Group, France, for the synthesis of exendin-4-VT750. They additionally thank Drs Isabelle Leclerc, Bryn Owen, and Aida Martinez-Sanchez for providing animal project licence access.

## Notes

### Competing Interest Statement

AT and BJ have received grant funding from Eli Lilly.

### Summary of Updates

New updated version

