## Supplementary figures and images for "Divergent acute *versus* prolonged pharmacological GLP-1R responses in adult beta cell-selective β-arrestin 2 knockout mice"

### Graphical Abstract

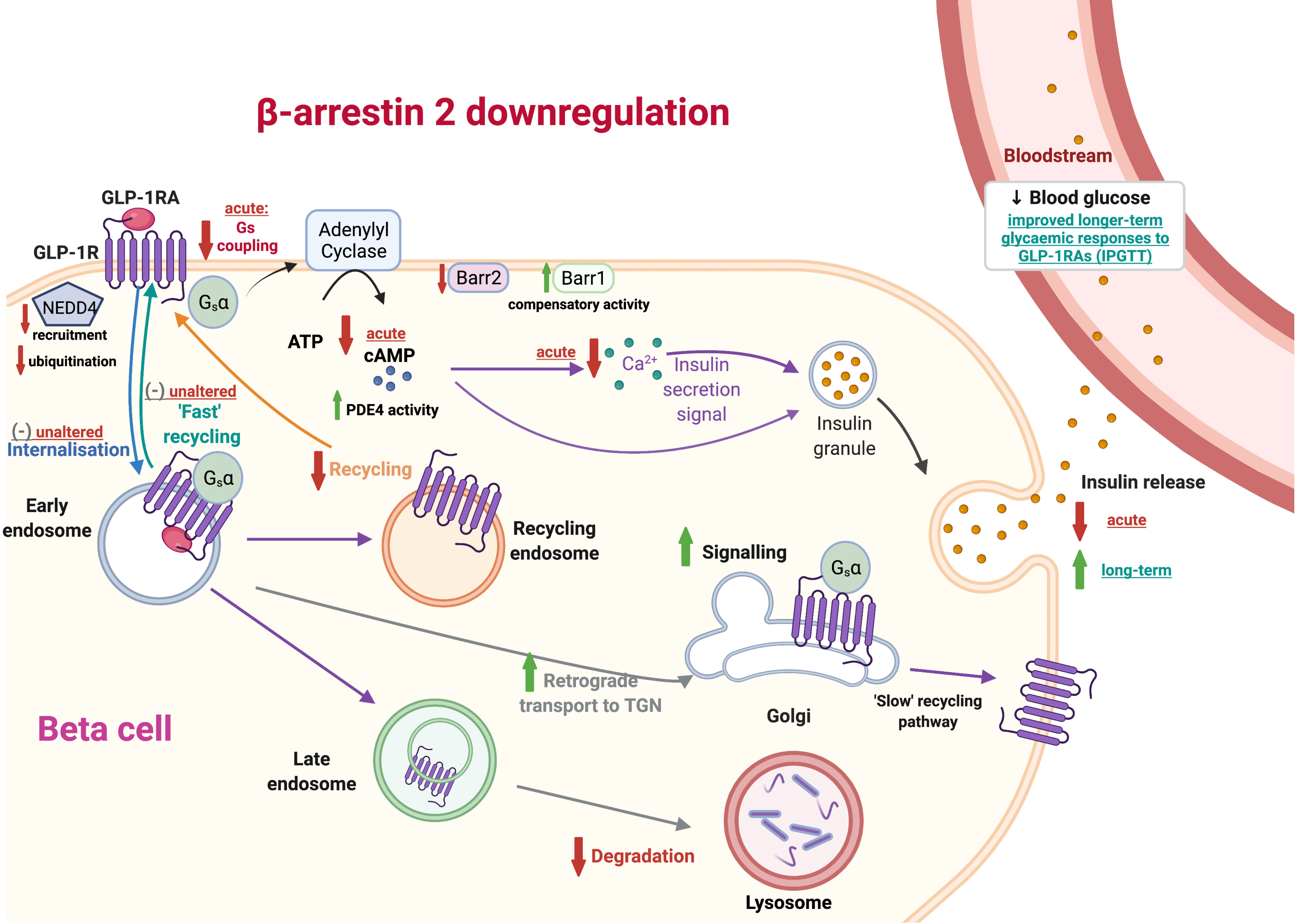
