## Supplemental Figures for "Divergent acute *versus* prolonged pharmacological GLP-1R responses in adult beta cell-selective β-arrestin 2 knockout mice"

### Supplemental Figure 1

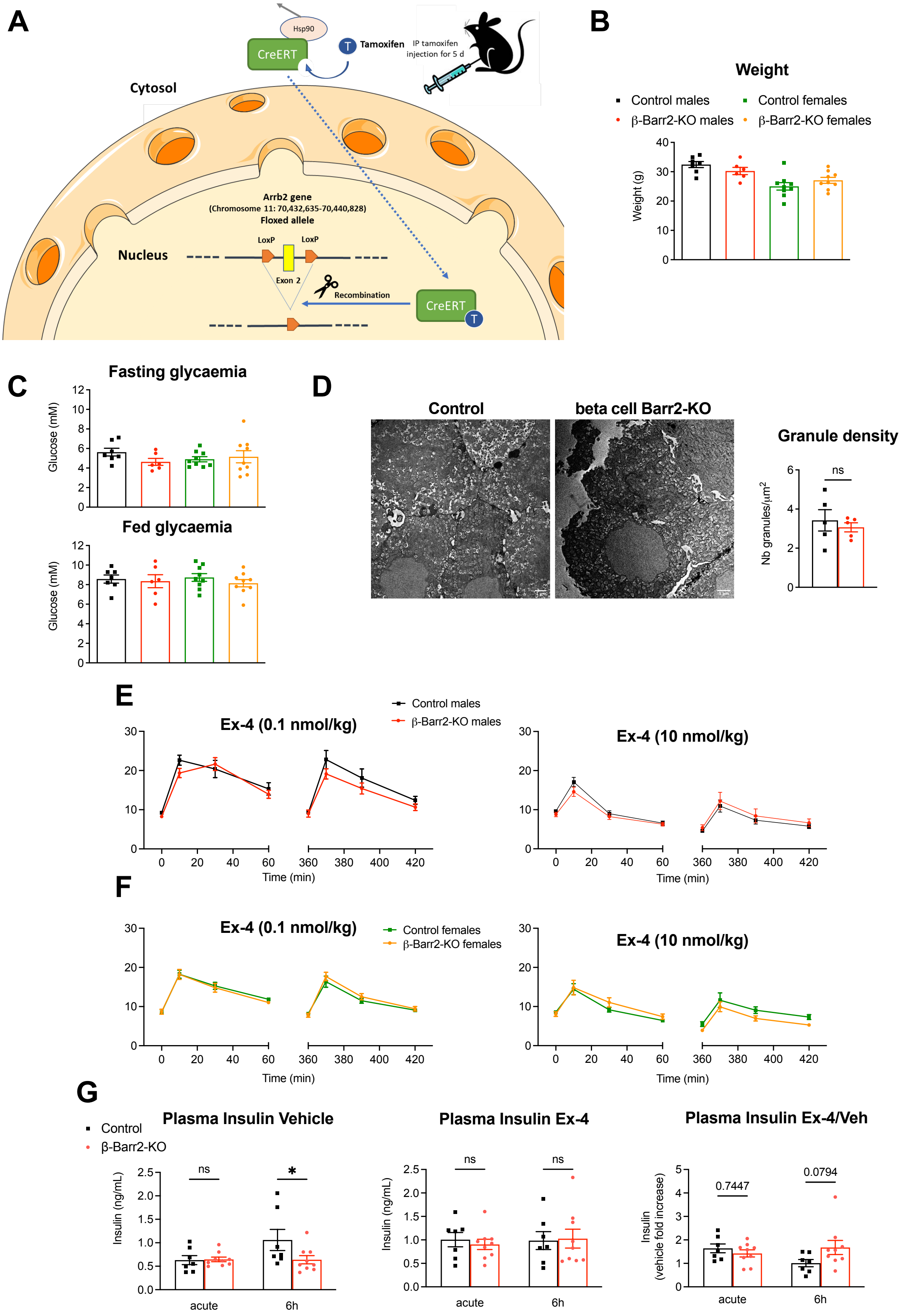

**Supplemental Figure 1. GLP-1R agonist responses in lean adult beta cell-selective β-arrestin 2 KO *vs* control mice – extra data.** IPGTTs (2 g/kg glucose i.p.) were performed concurrently with, or 6 h after, i.p. administration of agonists or vehicle (saline). (**A**) Diagram depicting the tamoxifen-induced adult beta cell-selective β-arrestin 2 KO mechanism using the Cre-lox system. (**B**, **C**) Weight (B), and fasting and fed glycaemia (C) of lean adult beta cell-selective β-arrestin 2 (β-Barr2) KO *vs* control male and female mice (n = 6-9 / genotype and sex, age: 20-24 weeks). (**D**) Representative EM images depicting islet ultrastructure and quantification of insulin granule densities in islets isolated from lean adult β-Barr2 KO *vs* control mice on chow diet (n = 5). (**E**) Glucose curves for vehicle or exendin-4 (Ex-4) administration at 0.1 and 10 nmol/kg in lean, male mice (n = 8 / genotype, age: 12-16 weeks). (**F**) Glucose curves for vehicle or Ex-4 administration at 0.1 and 10 nmol/kg in lean, female mice (n = 9 / genotype, age: 12-16 weeks). (**G**) Absolute and fold-change *vs* vehicle values for 10-min plasma insulin concentrations during IPGTTs (2 g/kg glucose i.p.) performed concurrently with, or 6 h after administration of vehicle or 1 nmol/kg Ex-4 in lean female mice (n = 7-9 / genotype, age: 12-16 weeks). Comparisons were performed using unpaired t-test or two-way ANOVA with Sidak's *post hoc* tests. Data are presented as mean ± SEM.

### Supplemental Figure 2

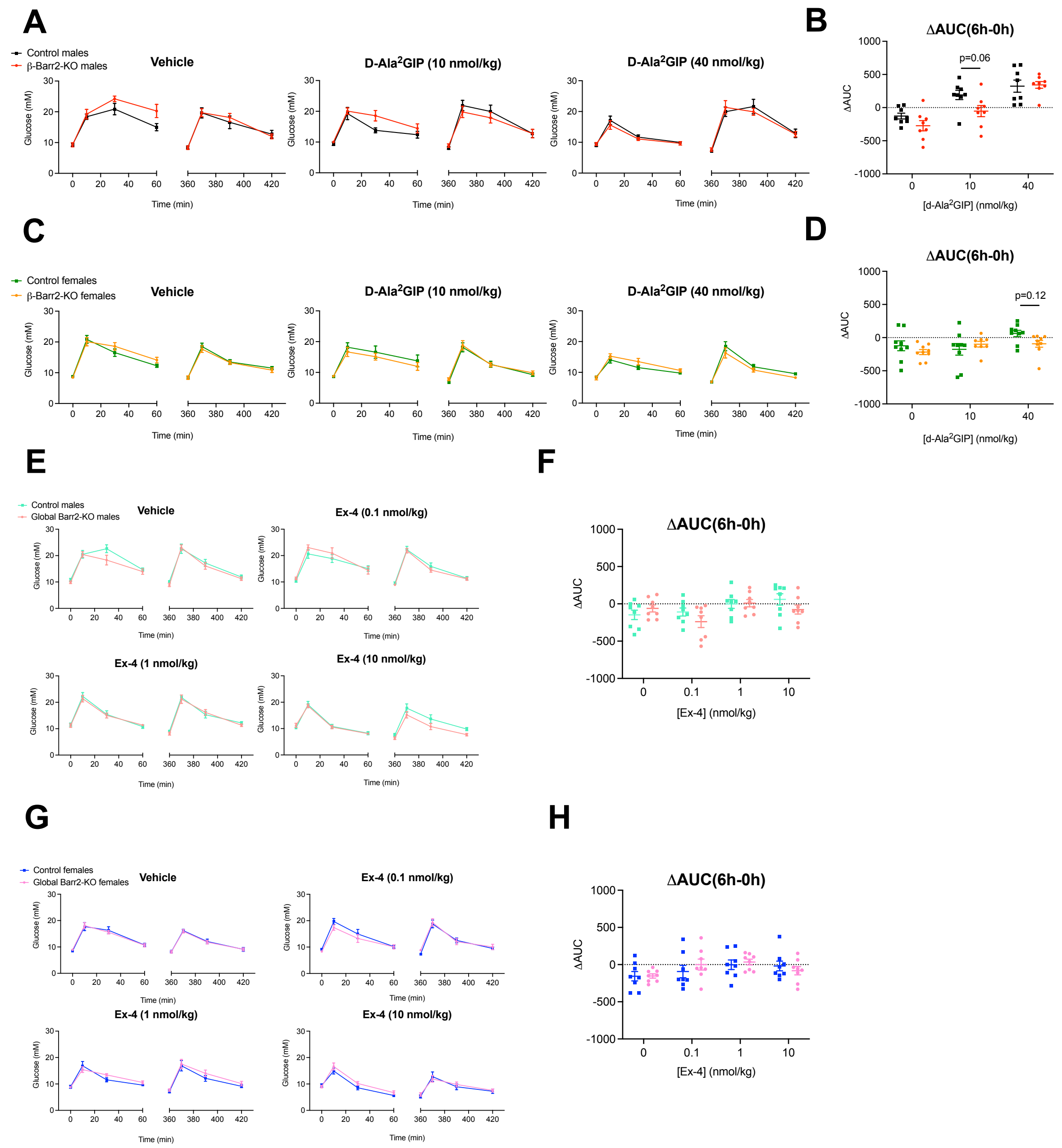


**Supplemental Figure 2. GIPR agonist responses in adult beta cell-selective β-arrestin 2 KO *vs* control mice and GLP-1R agonist responses in whole body β-arrestin 2 KO *vs* control mice.** IPGTTs (2 g/kg glucose i.p.) performed concurrently with or 6 h after i.p. administration of agonists or vehicle (saline). (**A**, **B**) Glucose curves (A) and corresponding ΔAUCs (6h-0h) (B) for vehicle or D-Ala$^2$-GIP administration at 10 and 40 nmol/kg in lean male mice (n = 8 / genotype, age: 18-26 weeks). (**C**, **D**) Glucose curves (C) and corresponding ΔAUCs (6h-0h) (D) for vehicle or D-Ala$^2$-GIP administration at 10 and 40 nmol/kg in lean female mice (n = 8-9 / genotype, age: 18-26 weeks). (**E**, **F**) Glucose curves (E) and corresponding ΔAUCs (6h-0h) (F) for vehicle or Ex-4 administration at 0.1, 1, and 10 nmol/kg in tamoxifen-inducible whole-body (R26-Cre-ERT) Barr2 KO and control lean males (n = 8 / genotype, age: 12-16 weeks). (**G**, **H**) Glucose curves (G) and corresponding ΔAUCs (6h-0h) (H) for vehicle or Ex-4 administration at 0.1, 1, and 10 nmol/kg in tamoxifen-inducible whole-body (R26-Cre-ERT) Barr2 KO and control lean females (n = 8 / genotype, age: 12-16 weeks). Comparisons were performed using two-way ANOVA or mixed-effects model with Sidak's *post hoc* tests. Data are presented as mean ± SEM.

### Supplemental Figure 3

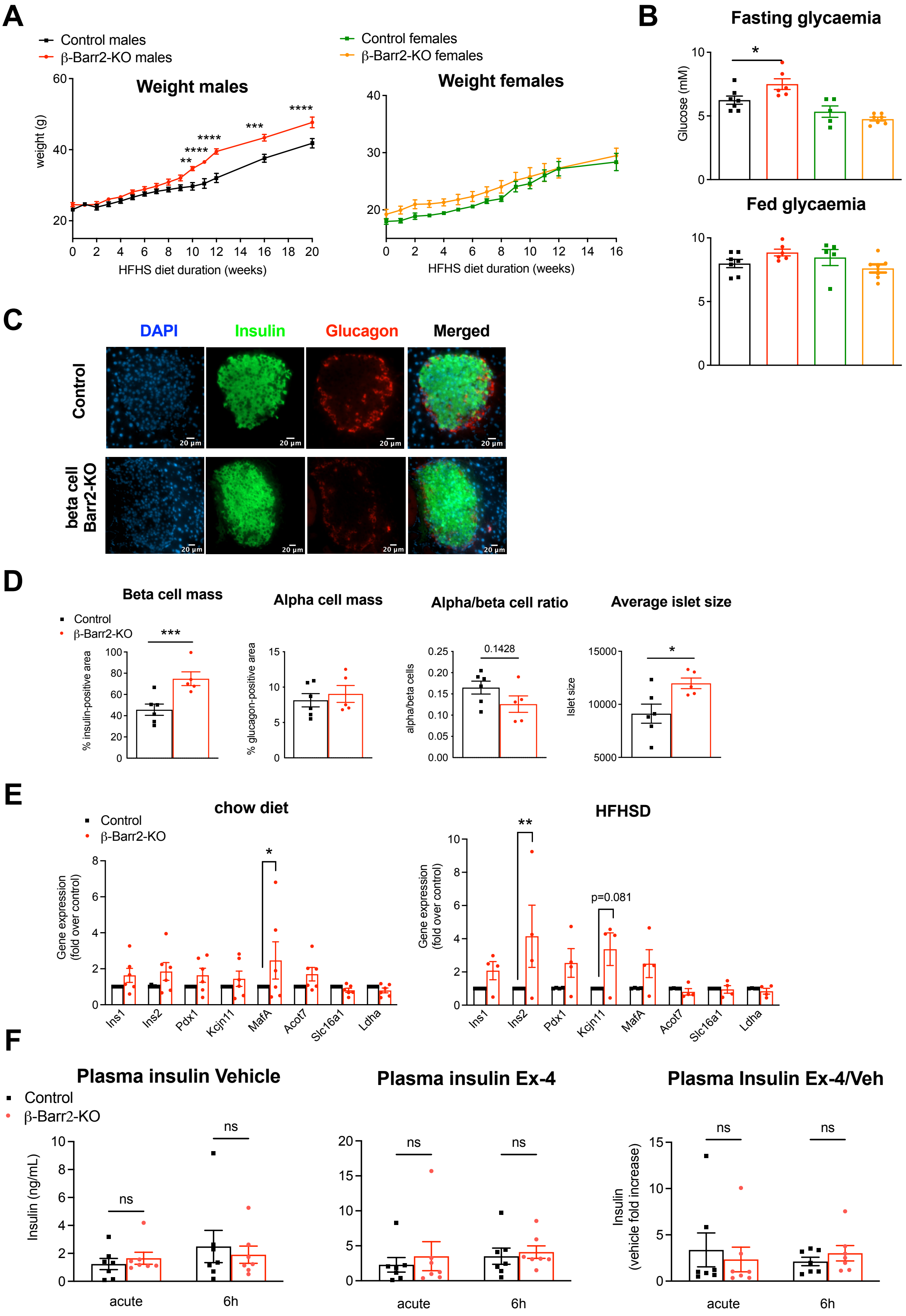

**Supplemental Figure 3. HFHS-fed adult beta cell-selective β-arrestin 2 KO *vs* control mice – extra data.** (**A**) Weekly weights of male and female mice after HFHS diet initiation (n = 7-9 / genotype and sex). (**B**) Fasting and fed glycaemia of HSHF diet-fed male and female mice (n = 5-7 / genotype and sex, duration of HSHF diet: 10-16 weeks). (**C**) Representative images of pancreatic sections from adult beta cell-selective β-arrestin 2 (β-Barr2) KO *vs* control mice on HFHS diet depicting islets with nuclei stained with DAPI (blue) and co-stained for insulin (green) and glucagon (red). (**D**) Quantifications of beta and alpha cell mass, alpha/beta cell mass ratio and average islet sizes in adult β-Barr2 KO *vs* control mice on HFHS diet (n = 5-6 / group). (**E**) Relative gene expression of selected beta cell-enriched and -disallowed genes in control *vs* adult β-Barr2 KO mice on chow (n = 6) or HFHS diet (n = 4). (**F**) Absolute and fold-change *vs* vehicle values for 10-min plasma insulin concentrations during IPGTTs (2 g/kg glucose i.p.) performed concurrently with or 6 h after administration of vehicle or 1 nmol/kg Ex-4 in mixed male and female mice on HFHS diet (n = 7 / genotype, duration of HFHS diet: 10-16 weeks). Comparisons were made using unpaired t-tests or one or two-way ANOVA with Sidak's *post hoc* tests. *p<0.05, **p<0.01, ***p<0.001, ****p<0.0001 *vs* control group. Data are presented as mean ± SEM.

### Supplemental Figure 4

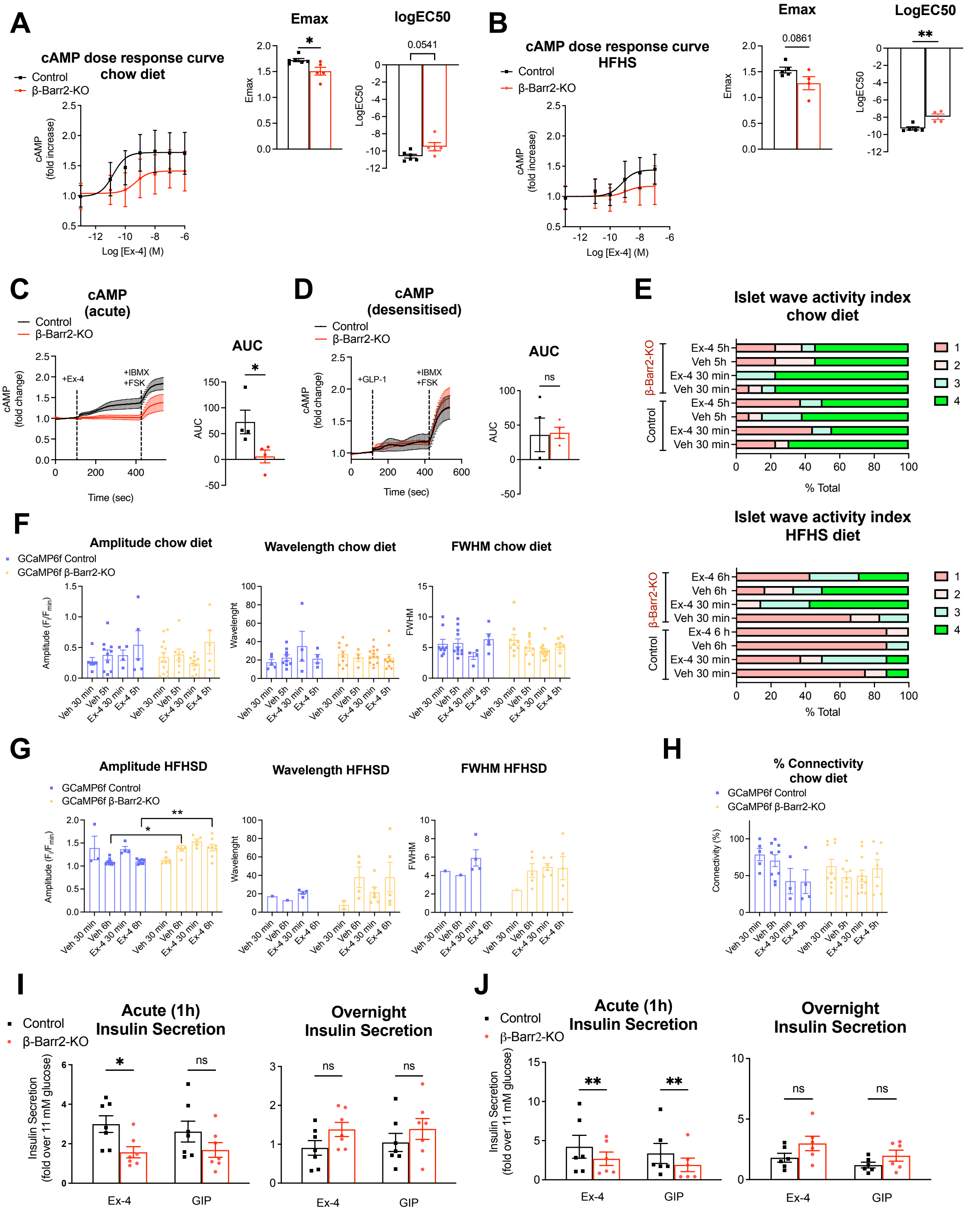


**Supplemental Figure 4. Extra *ex vivo* signalling data from adult beta cell-selective β-arrestin 2 KO *vs* control islets.** (**A**, **B**) cAMP dose-response curves and corresponding Emax and logEC50 values in dispersed islets isolated from adult beta cell-selective β-arrestin 2 (β-Barr2) KO and control mice on (A) chow diet (n= 5-6 / group) or (B) HFHS diet (n= 4-5 /

group). (**C**) cAMP responses over time from cADDis-infected beta cell-specific Barr2 KO *vs* control islets in response to 100 nM Ex-4 followed by 100 μM IBMX + 10 μM forskolin (FSK); AUCs calculated for the Ex-4 treatment period (n = 4 / genotype). (**D**) cAMP responses over time from cADDis-infected β-Barr2 KO *vs* control islets pre-treated with 1 nM Ex-4 for 16 h (overnight) in response to 1 nM GLP-1 stimulation followed by 100 μM IBMX + 10 μM FSK; AUCs calculated for the GLP-1 treatment period (n = 4 / genotype). (**E**) Percentage of islets within each wave activity index category for chow and HFHS diet animals (n = 6-13 islets / condition). (**F**, **G**) Calcium wave characteristics: amplitude, wavelength, and full width at half maximum (FWHM) for beta cell-selective β-arrestin 2 (β-Barr2) KO *vs* control GCaMP6f islets from donors implanted in the anterior chamber of the eye from (F) chow and (G) HFSH diet experiments (n = 0-13 islets / condition). (**H**) Percentage of connectivity for β-Barr2 KO *vs* control GCaMP6f islets from chow diet animals implanted in the anterior chamber of the eye of chow diet-fed WT animals, which received Ex-4 10 nmol/kg or saline (vehicle) i.p. (n = 2-11 islets / condition). (**I**, **J**) Insulin secretion assays from control *vs* β-Barr2 KO islets treated with 100 nM Ex-4 or 100 nM GIP (fold over 11 mM glucose vehicle levels) for 1 h (acute) or 16 h (overnight) isolated from mice on (I) chow diet (n = 8 / genotype) or (J) HFHS diet (n = 6 / genotype). Comparisons were made with t-tests or two-way ANOVA with Sidak's *post hoc* tests. *p<0.05, **p<0.01 *vs* control group. Data are presented as mean ± SEM.

### Supplemental Figure 5

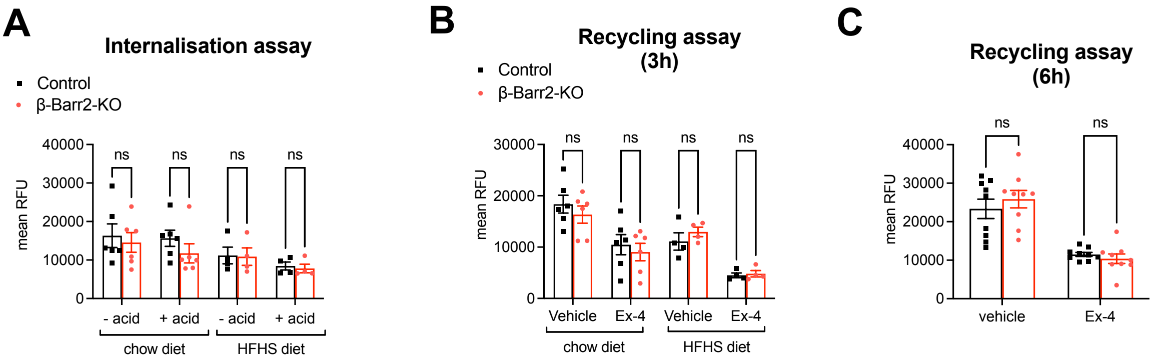


**Supplemental Figure 5. Extra *ex vivo* trafficking data from adult beta cell-selective β-arrestin 2 KO *vs* control islets.** (**A**) Ex-4-TMR fluorescence at 1 h in β-Barr2 KO *vs* control islets treated or not with acetic acid wash for 5 min before imaging for animals on chow (n = 6 / genotype) or HFHS diet (n = 4 / genotype). (**B**) Quantification of TMR fluorescence in β-Barr2 KO *vs* control islets pre-treated with vehicle or 100 nM Ex-4 for 1 h and then washed and treated with 100 nM Ex-4-TMR for 3 h for animals on chow (n = 6 / genotype) or HFHS diet (n = 4 / genotype). (**C**) TMR fluorescence in β-Barr2 KO *vs* control islets pre-treated with vehicle or 100 nM Ex-4 for 1 h and then washed and treated with 100 nM Ex-4-TMR for 6 h in chow diet animals (n = 9 / genotype). Comparisons were made with two-way ANOVA with Sidak's *post hoc* tests. Data are presented as mean ± SEM.

### Supplemental Figure 6

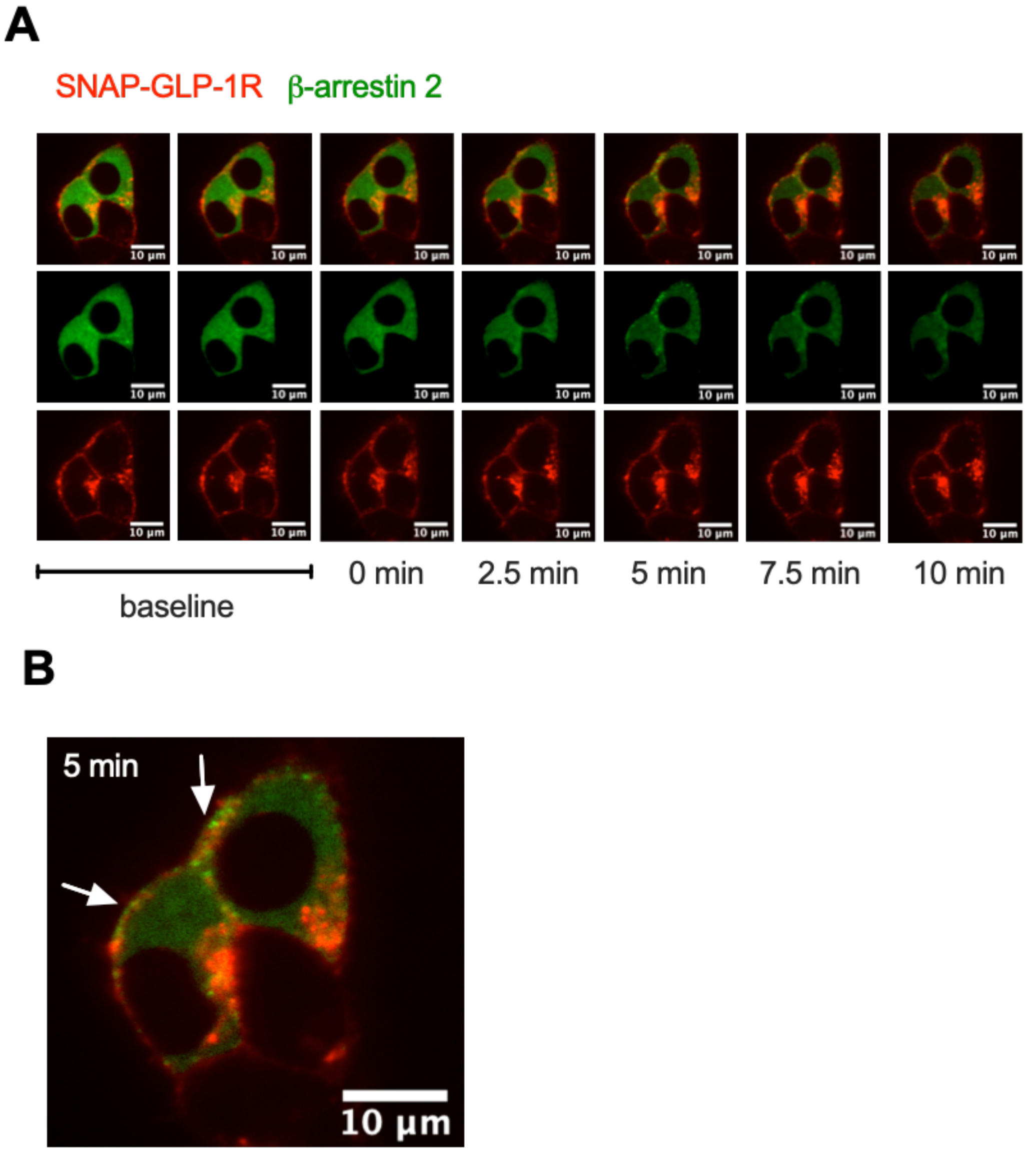


**Supplemental Figure 6. β-arrestin 2 recruitment dynamics in INS-1 832/3 SNAP-GLP-1R cells.** (**A**) Time-lapse spinning disk imaging of SNAP-GLP-1R (red) and β-arrestin 2-GFP (green) dynamic profiles in INS-1 832/3 SNAP-GLP-1R cells in response to 100 nM Ex-4. Selected time frames are depicted for the green, red, and merged channels. **(B**) High magnification image for the 5-min stimulation time-point from (A). Areas of SNAP-GLP-1R - β-arrestin 2-GFP co-localisation are indicated with white arrows.

#### Supplemental Figure 7

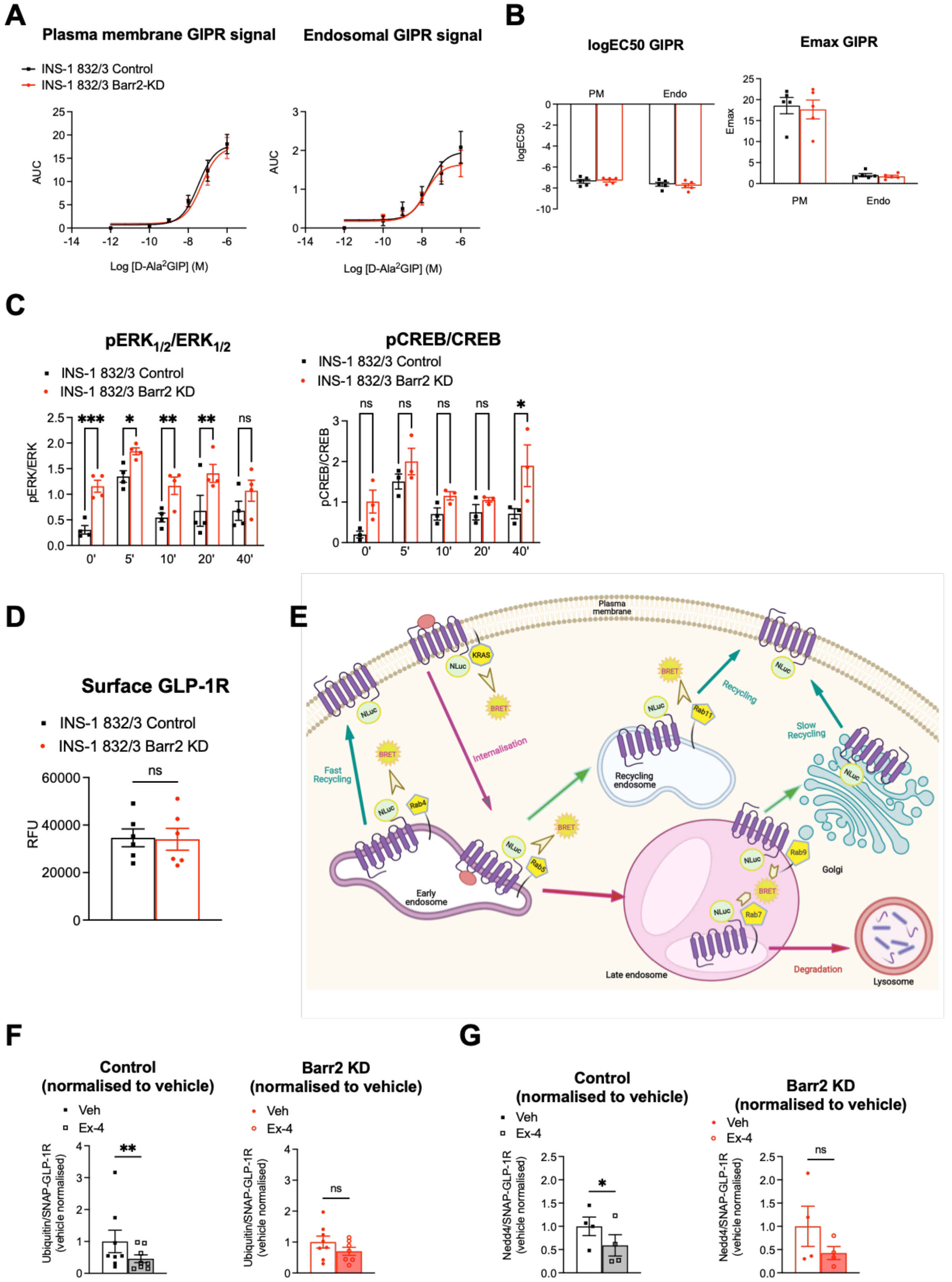


**Supplemental Figure 7. Extra data from β-arrestin 2 KD *vs* control INS-1 832/3 cell lines.**

(**A**) D-Ala$^2$GIP dose-response AUC curves for Nb37-SmBiT and CAAX-LgBiT (plasma membrane) or Endofin-LgBiT (endosomal) signal complementation assays (n = 5). (**B**) LogEC50 and Emax values calculated from (A). (**C**) Quantification of pERK1/2 over total ERK1/2, and pCREB over total CREB using densitometry analysis (n = 4 for pERK1/2 and n = 3 for pCREB). (**D**) Endogenous surface GLP-1R receptor levels in untreated β-arrestin 2 (Barr2) KD *vs* control INS-1 832/3 cells quantified by labelling for 1 h with 1 μM Ex-9-TMR (n = 6). (**E**) Schematic representation of the principle of NanoBRET-based subcellular localisation assays. (**F**) Quantification of SNAP-GLP-1R deubiquitination in response to 10 min of 100 nM Ex-4 exposure in INS-1 832/3 SNAP-GLP-1R control and Barr2 KD cells. Data from Figure 7F normalised to vehicle conditions for each cell type. (**G**) Quantification of loss of HA-NEDD4 - SNAP-GLP-1R interaction in response to 10 min of 100 nM Ex-4 exposure in INS-1 832/3 SNAP-GLP-1R Barr2 KD and control cells. Data from Figure 7G normalised to vehicle conditions for each cell type. Comparisons were made with ratio-t-tests and two-way ANOVA with Sidak's *post hoc* tests. *p<0.05, **p<0.01, ***p<0.001 *vs* control group. Data are presented as mean ± SEM.
